# Systematic evidence map of research on life-history traits in native and introduced earthworms

**DOI:** 10.64898/2026.09.18.752271

**Authors:** Karina Vanadzina, Kaushalya Nagahawatte, Deshan Rupasinghe, George G. Brown, Erin K. Cameron, Sylvain Gérard, Céline Pelosi, John W. Reynolds, Alyssa Rice, Helen R. P. Phillips

**Affiliations:** Faculty of Biological and Environmental Sciences, University of Helsinki, Helsinki, Finland; Universidade Federal do Paraná, Curitiba, Brazil; Embrapa Forestry, Colombo, Brazil; Department of Biology, Saint Mary’s University, Halifax, NS, Canada; INRAE, Avignon Université, UMR EMMAH, F-84000, Avignon, France; Oligochaetology Laboratory, 9-1250 Weber Street East, Kitchener, ON, Canada; New Brunswick Museum, Saint John, NB, Canada

**Keywords:** life-history traits, earthworms, invasive species, evidence map, ecotoxicity

## Abstract

Invasive species constitute one of the most significant threats to biodiversity worldwide, and are responsible for more than $400 billion in losses each year. An organism’s life-history strategy, defined by how it allocates resources among growth, survival, and reproduction, plays a central role in determining whether it becomes invasive. While the life-history profiles of aboveground invaders are increasingly well characterised, those of invasive species below ground remain poorly understood. Earthworms are a diverse group of soil invertebrates that can help fill this knowledge gap. Of the ∼5,800 earthworm species described to date, nearly 2% are cosmopolitan, meaning they have achieved wide geographic distributions despite limited ability for active dispersal. While life-history traits are likely to contribute to their worldwide distribution, the existing research on earthworm trait variation has not been systematically reviewed. To address this gap, we assessed the taxonomic scope, geographic coverage, and methodologies of studies on earthworm life history traits for species occurring in the US and Brazil, spanning a broad latitudinal range. Our literature search yielded over 8,000 articles across 180 species and revealed that introduced species have received considerably more research attention than native species. Research was concentrated on a few model species, most studies were experimental, and variation in life history traits was most commonly assessed in response to chemical compounds, vermicompost feed type, and environmental factors such as temperature, moisture and soil composition. Variation in adult survival was reported most frequently, followed by adult growth and cocoon production, with the bulk of life history research conducted in India (20%) and China (9%). To better understand the effect invasive earthworm species might have on native fauna, future research on life histories should address taxonomic biases and prioritise detailed study of the life cycles of native species in their natural habitat.

## I. INTRODUCTION

Invasive species, i.e., organisms that establish and spread in habitats outside of their native range, pose a considerable threat to biodiversity (Blackburn *et al*., 2014; Kumschick *et al*., 2015; Pyšek *et al*., 2020b) and the economy worldwide (Cuthbert *et al*., 2022; Bradshaw *et al*., 2024). Introductions of non-native organisms are occurring at unprecedented rates, with about 200 new introductions recorded worldwide each year (IPBES; Roy *et al*., 2023). This is primarily due to an intensification of global trade, agricultural activity and formation of new trade routes (Seebens *et al*., 2017). Other human-driven stressors such as climate change and habitat modification are expected to further exacerbate the effects of biological invasions by increasing the survival, spread and persistence of non-native species (Walther *et al*., 2009; Hulme, 2017).

Among introduced species, only a fraction establish and become invasive. Propagule pressure (also known as introduction effort) is a key factor that determines whether a population successfully establishes in a new habitat. Propagules may include adults, juveniles or other life stages, and propagule pressure reflects the number released (i.e., propagule size) and the frequency of release events (i.e., propagule number) (Lockwood, Cassey & Blackburn, 2005; Simberloff, 2009). Life-history (LH) traits characterising survival, development and reproduction of introduced species are often considered when examining the establishment success of invasive species (Sakai *et al*., 2001). These traits can affect both the number of propagules produced and the frequency with which they might be released. They can also determine whether founder populations not only grow beyond the vulnerable early stage, but also whether they subsequently spread (Blackburn, Lockwood & Cassey, 2015). For example, species with frequent breeding and large clutch sizes will require lower propagule pressure for establishment and will grow and spread more successfully than species with small number of young and a single reproductive episode. By linking species characteristics to invasion outcomes, trait-based approaches provide a predictive framework for identifying potentially invasive species (Sinclair *et al*., 2020; Pyšek *et al*., 2020a). These predictions can inform mitigation and management strategies that reduce ecological impacts and lower the direct and indirect economic costs associated with biological invasions (Cuthbert *et al*., 2025).

Recent decades have seen the development of large trait databases that have facilitated trait-based analyses, e.g., in plants (TRY; Kattge *et al*., (2011)), birds (AVONET; Tobias *et al*., (2022)), reptiles (RepTraits; Oskyrko *et al*., (2024)), fish (FishBase; Froese & Pauly (2025)), mammals (Jones *et al*., 2009), amphibians (Oliveira *et al*., 2017) and marine animals (McClain *et al*., 2025). The majority of trait-based studies of invasion potential have focused on plants (Hodgins, Bock & Rieseberg, 2018; Palma *et al*., 2021) and aboveground animals (Chown & McGeoch, 2023), even though ∼60% of global biodiversity, and a quarter of all animal species, is concentrated belowground (Decaëns *et al*., 2006; Anthony, Bender & van der Heijden, 2023). Large-scale studies using the traits of belowground fauna remain rare (e.g., Neyret *et al*., 2024; Lu *et al*., 2025), hindered by lack of global trait databases and data standardisation initiatives (Joimel *etal.*, 2024). Traits such as high fecundity and short reproductive cycles (in amphibians, reptiles and mammals; Capellini *et al*., 2015; Allen, Street & Capellini, 2017)) or repeated breeding attempts with low investment in each brood (in birds; Sol *et al*., 2012)) are important for aboveground taxa. However, as previous studies have revealed a geographical mismatch between the richness of above- and belowground organisms (Cameron *et al*., 2019; Phillips *et al*., 2019; Potapov *et al*., 2023), their trait composition and LH strategies are also likely to differ. Greater insight into the trait profiles of native and introduced belowground species is important for understanding the establishment of invasive species, particularly when combined with additional global change stressors. As species with similar traits are expected to respond similarly to global change (Bello *et al*., 2021; Green *et al*., 2022), trait analyses can reveal general patterns of vulnerability and resilience among belowground fauna.

Earthworms are a key group of soil macrofauna that are well suited for investigating invasion pathways belowground. They play a crucial role in ecosystem functioning (Blouin *et al*., 2013), have a long history of study (Brown *et al*., 2003), and are some of the largest soil invertebrates. Their ease of identification has promoted extensive research (Decaëns, 2010), resulting in a more detailed understanding of their traits compared to smaller groups. Around 120 (Blakemore, 2009) of the estimated 5,800 earthworm species (Misirlioğlu *et al*., 2023) are cosmopolitan (or ‘peregrine’ in field-specific literature). This means that they have attained large geographical distribution beyond their native range (Hendrix *et al*., 2008; Blakemore, 2009) despite their low rate of population expansion (1.4-14.1 m per year based on estimates from *Lumbricus terrestris* (Ligthart & Peek, 1997) and *Lumbricus rubellus* (Marinissen & van den Bosch, 1992) but can exceed 17.9 m in *Dendrobaena octaedra* (Cameron & Bayne, 2015)). The spread of cosmopolitan species is facilitated mainly through transport by humans, including through imported plants or historical use of soil as ship’s ballast, the abandonment of fishing bait and dispersal via roads (Hendrix & Bohlen, 2002; Cameron, Bayne & Clapperton, 2007; Hendrix *et al*., 2008) and waterways (Costello, Tiegs & Lamberti, 2011). However, it is the combination of suitable environmental factors and intrinsic traits that typically enables introduced species to establish and become invasive (Richardson & Pyšek, 2006). It is still unclear how many cosmopolitan species qualify as invasive, but introduced earthworms have had lasting effects on biodiversity and ecosystem processes in invaded habitats, as illustrated by the invasion of North American hardwood forests by Lumbricidae species from Europe (Baumann *et al*., 2024), and jumping worms, mainly *Amynthas* spp., from Asia (Chang *et al*., 2021). Another example is the global spread of *Pontoscolex corethrurus* throughout tropical regions (Chauvel *et al*., 1999; Römbke et al., 2009). These introduced species can dramatically transform soil structure and chemical properties (Chauvel *et al*., 1999; Eisenhauer *et al*., 2007), reshape microbial communities (Dempsey *et al*., 2013) and speed up removal of the litter layer, which in turn changes native understory vegetation (Craven *et al*., 2017) and reduces the abundance and richness of soil macroinvertebrate fauna (Demetrio *et al*., 2023) and other animals such as ground-nesting songbirds (Loss & Blair, 2014).

Research on invasive earthworm populations suggests that certain intrinsic traits of non-native earthworms, particularly those related to reproduction, may be central to their establishment and spread. A recent large-scale study of North American invasions indicates that parthenogenesis (i.e., capacity to reproduce without mating) is more widespread among invasive species compared to native ones, with the ability of a single individual to establish a population likely driving their invasiveness (Terhivuo & Saura, 2006; Mathieu *et al*., 2024). In line with observations in aboveground clades, invasive earthworms are often characterised by high reproductive output, short incubation periods and fast life cycles, such as the ‘boom and bust’ annual cycle of *Amynthas* jumping worms (Hendrix *et al*., 2008; Chang *et al*., 2021). The success of establishment in new, potentially harsh habitats, is also affected by the earthworm life stage. Earthworm cocoons (egg capsules) of invasive species have been reported to survive extreme cold (Holmstrup, 2003; Görres, Bellitürk & Melnichuk, 2016) and drought events (Holmstrup, 1995) as well as flooding (Plum, 2005). Adults of some species (e.g., *D. octaedra*) can tolerate freezing by rapid accumulation of glucose (Rasmussen & Holmstrup, 2002). Cocoons are typically deposited in shallow soil layers, where they persist through frost and drought via dehydration, losing nearly all osmotically active water and instead accumulating protective sugars and polyols, as documented in *Aporrectodea caliginosa*, *Allolobophora chlorotica*, and *D. octaedra* (Holmstrup & Westh, 1994, 1995). Although early reviews argued that traits poorly predict earthworm invasiveness because many are shared by both invasive and non-invasive species (Hendrix & Bohlen, 2002; Hendrix *et al*., 2008), a systematic comparative study of trait profiles in invasive versus non-invasive earthworms has yet to be conducted. This would facilitate the understanding on how the intrinsic traits of the invading species affect their establishment success.

Here we applied a systematic mapping framework (James, Randall & Haddaway, 2016) to identify knowledge clusters and gaps in earthworm LH trait research, using a sample of ∼500 native and introduced species from the US and Brazil as our basis for extrapolating global trends. We also aimed to quantify taxonomic biases in the literature, as there are a number of patterns we expected to see within the literature. Firstly, we anticipated dominance of model species due to the use of LH traits as important endpoints in ecotoxicity and waste management studies. The OECD toxicity test guidelines – Tests 207 and 222 (OECD, 1984, 2016) – recommend the use of highly fecund model species with short life-cycles such as *Eisenia fetida* and *Eisenia andrei* [Note: historically these two species were considered synonymous, thus older literature often do not distinguish between them], and vermicomposting studies employ species with high capacity for feeding and biodegradation (Sinha *et al*., 2002). Secondly, similar to other synthesis approaches in ecology (Martin, Blossey & Ellis, 2012), we expected to observe an over-representation of studies from temperate regions, particularly Europe and North America, with less studies from tropical habitats (Cameron *et al*., 2018). Hence, we had the following main objectives:

1. Assess the distribution of research effort among species and LH traits;
2. Identify what study design (laboratory vs field; experimental vs observational) is most represented among articles investigating LH variation;
3. Identify biases in the geographic distribution of research effort.

With this systematic literature analysis, we intend to provide key recommendations to guide future research into earthworm LH trait variation and its potential drivers, particularly with respect to species with high invasion potential.

## II. MATERIALS AND METHODS

### (1) Search methods

Our systematic map reviews the evidence base of experimental and observational studies on LH trait variation in earthworms. We focused on native and introduced species in the United States and Brazil, sourced from the GloWorm database (Phillips & Cameron, 2023). By focusing on these two countries, the data provide a broad latitudinal distribution, including a mix of tropical and temperate species, from two countries with a history of earthworm research and evidence of earthworm invasions. The species list contained 147 species documented in the US, 292 in Brazil, 28 present in both countries and 82 synonyms. Each species was categorised as native or introduced to the country, with the likely continent of origin provided for introduced species (Supplementary Material: Data S1). The study methods follow existing guidelines (O’Dea *et al*., 2021; Foo *et al*., 2021) and our preregistration (Vanadzina *et al*., 2025), which was submitted before the article screening stage. To ensure transparency, deviations from the preregistration have been highlighted below.

The finalised search strings were run on 17.11.2025 using Web of Science (WoS) Expanded and Scopus API interfaces in R Project and included all available records up to 17.11.2025, with the earliest one dating back to 1926 (full search strings in SM: Methods S1). To obtain research effort associated with each species, the search string was constructed as “species name” and terms relating to life history including early-life stages, reproduction, growth, metabolism, lifespan and mortality. As detailed in the preregistration, the output from a search string on earthworm reproduction was used to identify the initial set of terms that repeated in the title and abstract of articles at least 5 times using the Rapid Automatic Keyword Extraction (RAKE) algorithm in the *litsearchr* R package (Grames *et al*., 2019). The terms were analysed as a network, and a threshold corresponding to 80% of the cumulative network strength was used to retain only the more strongly connected terms in the final sample. For the final search string, we consulted T-SITA (Pey *et al*., 2014), Ecotaxonomy (Sandmann, Scheu & Potapov, 2019), Edwards and Arancon (2022), and nematode LH traits in Zhang *et al*. (2024). The search strategy was designed to be both sensitive, covering a broad range of LH traits, and specific, minimising the overlap with irrelevant terms. The search yielded 12,050 and 6,523 records on WoS and Scopus, respectively, after which duplicated entries were removed using *litsearchr* package resulting in 8,333 articles (see SM: Fig. S1 for PRISMA chart).

### (2) Article screening

The screening steps for both initial and full-text screening were carried out as described in the preregistration (SM: Fig. S1). The articles were sorted by relevance based on a score that reflected the number of relevant keywords in the abstracts and titles. This allowed us to reject articles with low relevancy from the initial screening and fast-track the full-text screening of high-relevancy articles (see SM: Methods S2 for keywords used). The 8,333 articles were further sorted into high-scorers (equal or more than 8 points, very likely to be relevant; 1356 articles), medium scorers (1-7 points, maybe relevant, 4962 articles), and 0 scorers (0 points, not= relevant; 2015 articles) depending on the number of relevant keywords in title and abstract using *stringr* R package (Wickham, 2023). In the keyword assignment list, keywords describing specific LH traits, e.g., cocoon production and hatching success, were assigned a higher value than more generic keywords such as ‘life-cycle’ or ‘reproduce’.

We used the Abstrackr software (https://abstrackr.com/home) for initial abstract and title screening primarily by observer DR who underwent training by observer KV to achieve inter-reviewer agreement of >95%; articles marked as ‘uncertain’ by observer DR were resolved by KV throughout the screening process. Abstrackr uses a machine learning algorithm that assigns a priority score to each article on a weekly basis. In our sample, screening was stopped at 0.1 after which 10 articles from each percentage point of priority score (0.09, 0.08 etc.) were screened at random; all articles in the random sample were rejections. The title and abstract screening was completed on 8 January 2026 yielding 2,296 articles suitable for full-text screening.

Following the title and abstract screening, we deviated from the pre-registration protocol as follows:

1) The screening process revealed that a proportion of toxicity articles rejected due to a 0 score could be relevant. An additional search containing terms ‘*mass, toxic, lethal, weight, surviv* (less restrictive spelling)’ were carried out on 0-scoring articles, yielding 828 additional articles to be screened in Abstrackr.
2) As KCL (Korean Journals Database) and SciELO (Scientific Electronic Library Online, focus on Latin America) databases were not included in the institutional subscription to the WoS, the search procedure was repeated for LH search strings and 1) all peregrine species for KCL and 2) all Brazil species for SciELO. This yielded a further 64 articles to be screened in Abstrackr.
3) The full-text screening of high-scoring articles was completed first, and we established that 51% of these articles focused exclusively on *E. fetida* or *E. andrei*, which are well-established model species in toxicology. Our initial search had yielded records of articles associated with these species; we extracted the IDs of all articles associated with *E. fetida* and/or *E. andrei* that did not appear in the search results of any other species. Before the full-text screening of 2,296 medium scorers, they were sorted into articles focusing (1) on *E. fetida* or *E. andrei* only (‘E’) or (2) on other species (‘NE’). This resulted in a 52% (E) vs. 48% (NE) split. Given that the split was similar to that among high-scoring articles, with a strong bias towards *Eisenia* spp., a random sample was expected to capture the main trends in the literature while reducing the manual coding burden. A combined manual and automated pdf search in Zotero referencing software (Corporation for Digital Scholarship, 2023) identified 2,029 articles with available pdfs, from which we then randomly selected 500 articles (250 E and 250 NE articles), representing a quarter of all medium scorers. We acknowledge that rare species, treatments and traits may be underestimated in this sample.

### (3) Data extraction

The following information was extracted from articles that were included in the full-text screening:

(1) **basic article meta-data:** keywords assigned by the authors (if available), country of the institution where research took place (or the institution of the first author if not specified), year of publication, further article details (author list, publication name, DOI, volume, issue);
(2) **study design:** laboratory or field experimental study (i.e., conditions were manipulated during the study), laboratory or field observational study (i.e., conditions were recorded but not manipulated during the study), review article, general species description (overview of species’ life history, typically presented as part of a taxonomic description), modelling study (mathematical modelling study incorporating data from experimental or field studies); binomial name of species or genus under study (as listed in the source);
(3) presence or absence of a set of **treatment types** for **experimental** laboratory or field studies that were varied or manipulated for the purposes of the experiment (i.e., independent variables), see Table 1 for description;
(4) presence or absence of a set of **LH traits** measured or observed as outcomes of field or laboratory studies (i.e., dependent variables), grouped into traits relating to offspring, survival, growth, timing (i.e., phenology) and others; see Table 2 for description. We recorded if LH traits of immature (cocoons, hatchlings, juveniles) or mature (adult) stages of earthworm life cycle were reported. Numbers of immature worms, as well as growth and survival rates for adults and immatures, were noted only if they characterised the whole experimental population. Generally, we focused on LH traits measured as part of repeated sampling designs, e.g., experimental groups or time points. The exception was juvenile:adult ratio, which was recorded only for independent population-level samples.

**Table 1.** Definitions of treatments used in experimental laboratory and field studies.

| Main treatment group | Treatment code | Description |
| --- | --- | --- |
| **Environment** | **Temp** | temperature treatments (could be soil/air) |
|  | **ColdHeatResistance** | resistance to extreme cold or heat specifically tested |
|  | **SoilpH** | soil pH varied |
|  | **SoilContent** | information on soil content, such as soil texture, e.g. silt, clay, and components, e.g. organic C content, reported |
|  | **SoilProfile** | different soils tested (all aspects characterising soil) |
|  | **SoilWater** | moisture content and/or water potential in soil varied |
|  | **Feed** | type or amount of feed varied, typically plant or tree leaf litter [excludes vermicomposting studies] |
| **Waste management**<br>Studies focus on earthworms as active agents of breaking down waste | **Vermicomposting** | studies using earthworms to break down different waste materials into nutrient-rich compost |
|  | **Bioremediation** | studies using earthworms to help clean contaminated soil by breaking down harmful substances |
|  | **Source & type of waste** |  |
|  | **Animal_Manure** | livestock/farm animal waste (e.g., manure, dung, slurry from cattle, poultry, pigs etc.) |
|  | **Animal_OtherByproduct** | animal produce other than manure (e.g., bonemeal, skin) |
|  | **Plant_Agri** | residues from agricultural plants/crops, e.g., corn leaves and other residues, fruit, straw, husks, bagasse [includes mushroom byproducts] |
|  | **Plant_Other** | other plant residues, e.g., tree litter, weeds, grass clippings |
|  | **Industrial_Sludge** | source of feed from factories or production plants (e.g., sugar cane mill, tannery), semi-solid organic residue generated during wastewater or sewage treatment |
|  | **Industrial_OtherByproduct** | source of feed from factories or production plants other than sludge (e.g., filter cake from sugar cane mill, tannery effluent) |
|  | **Municipal_Sludge** | semi-solid organic residue from sewage facilities or urban waste waters |
|  | **Municipal_OtherByproduct** | residue from byproducts originating from sewage facilities or urban wastewaters (e.g., wastewater effluent) |
|  | **Domestic_Commercial** | organic waste from residential dwellings, canteens, marketplace, shops (e.g., peels, compost, kitchen waste) |
|  | **Paper_Wood** | cardboard, paper, wood chips or sawdust particles used as bulking agents |
|  | **Other_Waste** | other types of waste not included above |
| **Toxicity**<br>Studies focus on earthworm response to chemical and physical toxicants | **HeavyMetals** | metallic element, typically with high atomic weight and density, such as cadmium, mercury lead, zinc, arsenic, iron, copper, frequently toxic to organisms |
|  | **Toxin** | toxin derived from an animal, plant or microbes |
|  | **Medicines** | veterinary or human pharmaceutical drugs; can also include naturally-derived medicines such as essential oils |
|  | **Pesticides** | all types of pesticides (incl. insecticides, herbicides, fungicides, nematicides, molluscicides) |
|  | **OPol** | carbon-based chemical pollutants (incl. DDT, dioxins, PCBs, PAHs, PFAS, VOCs) |
|  | **SoilAmendments** | material added to the soil to improve its properties and enhance plant growth (biochar, vermiculite, limestone, biosolids, i.e., stabilised and treated sewage sludge) |
|  | **Fertiliser** | plant fertiliser manufactured in an industrial process, e.g., nitrogen, phosphorus, and potassium |
|  | **Salinity** | different levels of salinity as treatment |
|  | **Plastics_Micro_Nano** | plastics/micro- or nano-plastics |
|  | **Nanoparticles** | nanoparticles as treatment (usually in combination with heavy metals) |
|  | **Radiation** | exposure to radiation treatments tested, e.g., UV-C rays or gamma |
|  | **ToxOther** | other type of toxicity treatment |
| **Interactions** | **Density** | the effect of density tested (variation in number/quantity of earthworms per group tested); only applies to the same species |
|  | **InterSpeciesInt** | the effect of species composition tested (if different species tested together) |
| **Ungrouped** | **Genetic** | test groups differ by genetic factors |
|  | **Land_use** | different land management treatments, e.g., presence or absence of tillage (i.e., agricultural preparation of soil), conventional vs. organic farming, fire |
|  | **Bacterial_culture** | test groups differ by bacterial culture added |
|  | **Other** | other treatment type not included above |

**Table 2.** Definitions of LH traits subject to treatments/observations in full-text papers.

| LH trait type | Variable | Description |
| --- | --- | --- |
| **Offspring traits: Cocoon** | **C_Length_dm** | cocoon size reported (could be provided as length, breadth, diameter or axis, typically in mm) |
|  | **C_Weight** | cocoon weight reported (could be for single or many, typically in mg) |
|  | **C_Nr/Prod_Rate** | total cocoon number or count or cocoon production rate reported (e.g., cocoons per worm per year); could also be described as reproductive rate, fecundity, fertility etc. |
|  | **C_Characteristics** | cocoon colour, shape or ornaments reported |
|  | **C_Photo_Drawing** | cocoon photo or drawing present |
|  | **C_Other** | other information relating to cocoon reported (e.g., depth found) |
| **Offspring traits: Hatchling & Juvenile** | **H_Length_dm / J_Length_dm** | hatchling/juvenile size reported (e.g., as length or diameter) |
|  | **H_Weight / J_Weight** | hatchling/juvenile weight |
|  | **H_Nr_per cocoon** | number of hatchlings per cocoon reported |
|  | **H_Prod / J_Prod** | total number or count of hatchlings/juveniles or hatchling/juvenile production rate per entity (e.g., 'per worm') and/or frequencies (e.g., 'per month', 'per year') reported |
|  | **J_Length_dm** | juvenile size reported (e.g., as length or diameter) |
|  | **J_Weight** | juvenile weight (could be single or many) |
|  | **J_Nr / Prod_Rate** | total number or count of juveniles or juvenile production rate per entity (e.g., 'per worm') and/or frequencies (e.g., 'per month', 'per year') reported |
|  | **H_Success_Rate** | proportion of cocoons yielding hatchlings (typically in %) |
|  | **HJ_Other** | other traits relating to hatchlings or juveniles |
| **Growth** | **G_IM_Tot** | growth (i.e., increase or decrease in body size expressed as mass, weight, length) in immature stages (i.e., hatchlings, juveniles, subadults, young) given |
|  | **G_A_Tot** | growth (i.e., increase or decrease in body size expressed as mass, weight, length) in mature/adult/clitellate stages given |
|  | **G_NS** | not specified if growth measured in adults or immatures |
|  | **IGR** | Instantaneous Growth Rate (IGR), i.e., growth at a specific moment in time reported |
| **Survival** | **S_IM_Tot** | mortality, survival, senescence or death rate of immature stages (i.e., hatchlings, juveniles, subadults, young) given |
|  | **S_A_Tot** | mortality, survival, senescence or death rate of adult/mature/clitellate stages reported |
|  | **S_NS** | mortality, survival, senescence or death rate reported but life stage not specified |
| **Timing** | **Incubation** | information relating to cocoon incubation period (also as developmental period) is present, e.g., duration in days |
|  | **Sex_Maturity** | information relating to sexual maturity is present (e.g., time until sexual maturity, emergence of clitellate worms, proportion (%) of sexually mature worms given or weight at maturity given) |
|  | **Start_Cocoon_Prod** | start of cocoon production period given |
|  | **End_Cocoon_Prod** | end of cocoon production period given |
|  | **Breed_Freq** | information on whether the breeding is continuous or discrete (i.e., has distinct breeding episodes) |
|  | **Breed_Period** | information on the length of breeding period annually (e.g., July and August each year) or per worm's lifespan (e.g., 200 days out of 600) |
|  | **Longevity** | information relating to the total length of life cycle, lifespan or longevity of the worm (e.g., from cocoon stage until death) |
|  | **Diapause** | information relating to diapause or quiescence (a response either to adverse environmental conditions or a normal part of life-cycle; earthworm stops feeding and becomes inactive); can apply to both adults and cocoons |
| **Other** | **Juvenile:adult_Ratio** | proportion between adults and immature stages in a population (i.e., cocoons, juveniles, hatchlings, subadults, young) reported; provides an estimate of reproductive potential and demographic structure within a sampled population. As a population-level variable, can be reported as comparisons between different populations or time points. |
|  | **Repro_Mode** | study focuses on variation in reproductive mode, e.g., proportion of parthenogenetic or amphimictic morphs in a population, biparental vs uniparental reproduction |
|  | **Egg** | reproductive variables relating to eggs or egg production (oogenesis) are present (size, number, count, volume) |
|  | **Sperm** | reproductive variables relating to sperm or sperm production (spermogenesis) are present (size, number, count, volume) |
|  | **Metabolic_Rate** | direct measurements of metabolic rate included (via oxygen consumption, carbon dioxide production or calorimetry, i.e., heat produced) |
|  | **Parental_Care** | information on parental care behaviours reported |

### (4) Data synthesis

All analyses and visualisations were performed in the R environment (R version 4.4.2; RStudio version 2026.06.0) (R Core Team, 2020; Posit team, 2026). We used *tidyverse* (Wickham *et al*., 2019) for data management and cleaning*, biscale* (Prener *et al*., 2025) and *sf* for visualisation of geographic distribution of included studies (Pebesma & Bivand, 2023), *ggplot2* for visualisation of bar charts, *ggsankey* for Sankey plot (Sjoberg, 2026), *ggupset* for UpSet plots (Ahlmann-Eltze, 2025). Package *bibliometrix* (Aria & Cuccurullo, 2017) was used for the analysis of keywords with the Fruchterman-Reingold algorithm for network layout and the spinglass algorithm to partition the network into clusters; keywords inside the same cluster are strongly connected while connections between clusters are weaker.

## III. RESULTS

### (1) Taxonomic distribution of research effort

Article counts associated with native and non-native earthworm species revealed strong disparities in research effort from articles included in the full-text screening (Fig. 1, see SM: Fig. S2 for research effort per species from the original search, *n* = 8,333). In the articles included after the full-text screening (*n* = 1,546, Fig. 1a, SM: Data S2), data on LH traits were recovered for a total of 79 species present in the US and Brazil (SM: Data S1). Articles on introduced species dominated, with a small proportion of all articles focusing on species native to these countries (Fig. 1b). The contrast was particularly striking for Brazil: although 82% of its earthworm species are native and the country has more earthworm species than the US, species native to Brazil were the focus of only 0.1% of all LH-trait articles analysed in our study. The introduced species were widespread cosmopolitans, primarily from the Lumbricidae family, with all except one (*Bimastos rubidus*), native to Europe. The other species belonged to the family Megascolecidae originally from Asia, Acanthodrilidae (*Microscolex dubius,* likely native to South America), Eudrilidae (*Eudrilus eugeniae*) and Benhamiidae (*Dichogaster bolaui*) native to Africa, and Rhinodrilidae (*P. corethrurus*) native to the neotropics. The standard ecotoxicology and composting species *E. fetida* and *E. andrei* were together associated with 44% of all articles returned in the search (SM: Data S1) meaning that these model species have received almost the same level of research attention as all other species taken together.

**Figure 1.**
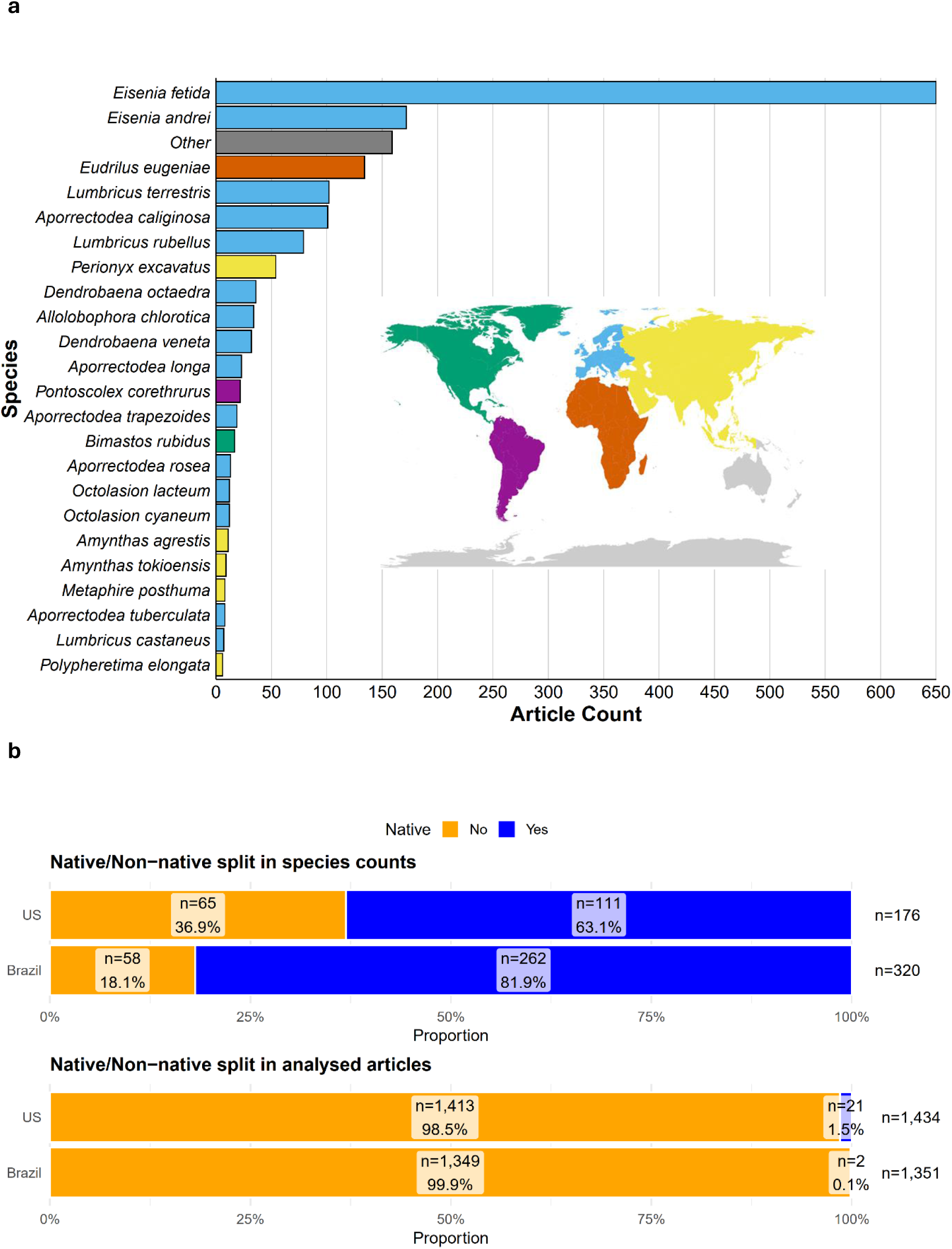
(a) Research effort per species based on articles included in the full-text screening (*n* = 1,546, only species with more than 5 articles displayed), colours indicate the species’ continent of geographical origin. ‘Other’ indicates species that appear in the literature but have been recorded in neither US or Brazil; (b) distribution of native/nonnative species occurring in the US and Brazil as counts (above) and as counts of articles associated with specific native/non-native species name in full-text screening.

### (2) Life history traits

Adult survival was the most commonly reported trait in LH literature (*n* = 829) followed by adult growth and cocoon production, with variation in these traits measured in response to treatment or observed under natural conditions (Fig. 2a; SM: Table S1). Adult survival is also frequently co-reported with cocoon production and adult growth (Fig. 2b). Information on survival and growth of immature life stages was included in only 25% of all reports on earthworm survival and growth trends. Cocoon production was the most commonly reported offspring trait (*n* = 656); by contrast, variation in egg and sperm traits was rarely investigated (*n* = 25). Time to sexual maturity was the most common phenological trait reported (*n* = 215). Some traits, such as variation in metabolic rate (i.e., energy consumption rate), longevity and reproductive mode were rarely investigated (all *n* <= 25 studies).

**Figure 2.**
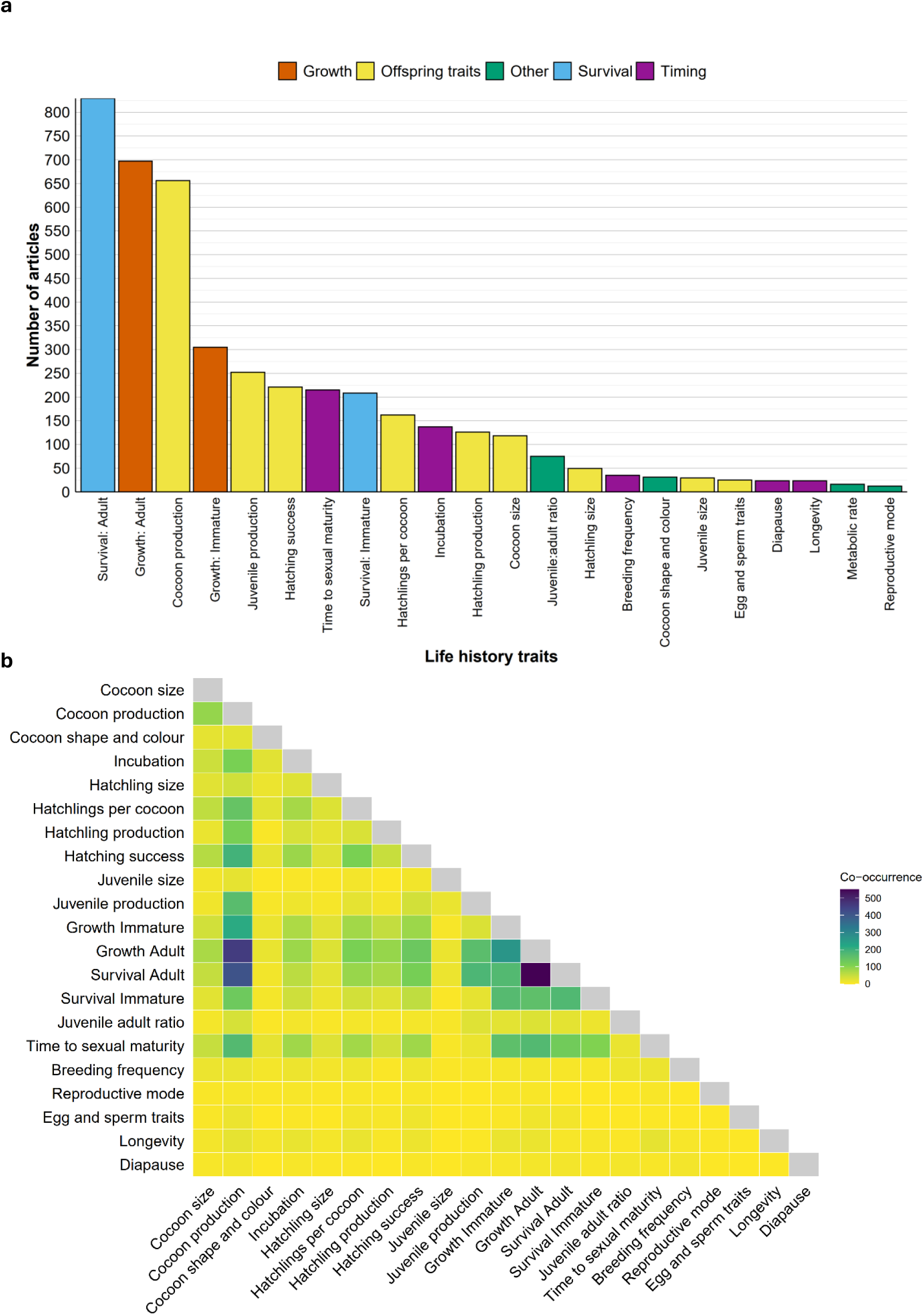
(a) Distribution and (b) heatmap of co-occurrence (per article) of life-history traits reported as response variables in articles included after full-text screening.

### (3) Study design and characteristics

The vast majority of studies investigating LH traits were experimental laboratory studies (85%), remotely followed by observational laboratory studies (3%) and experimental field studies (3%) (Fig. 3; SM: Table S2). If an article combined different study designs, it was most frequently an experimental study carried out in both laboratory or in the field, or a lab-based experiment with an observational field component. Studies examining earthworm response to chemical or physical toxicants represented the most common treatment type, followed by waste management and variation in environmental factors (Fig. 4a; SM: Table S3). A number of studies investigated combined treatment types, particularly toxicity treatments and environmental factors or waste management (Fig. 4a). Adult survival, adult growth and cocoon production were the most commonly reported LH traits across studies examining the three main treatment types (Fig. 4b). The co-occurrence network of keywords associated with articles included in the full-text screening further supported this finding. It highlights distinct clusters of articles on pollutants, pesticides, waste management and environmental factors (Fig. 5a). The number of studies published each year showed a sharp increase after the mid-1980s. However, in contrast to studies on waste management and ecotoxicity which were increasing exponentially, the number of studies assessing the effect of environment on LH traits remained at the 2005-level (Fig. 5b). A further breakdown of the treatment groups by subcategory indicated that toxicity treatments were dominated by the assessment of heavy metals, pesticides, and organic pollutants, with heavy metal treatments often combined with soil-profile comparisons, while waste-management studies predominantly focused on comparing animal waste, remains from agricultural plants and industrial sludge as substrates for vermicomposting (Fig. 6; SM: Table S4).

**Figure 3.**
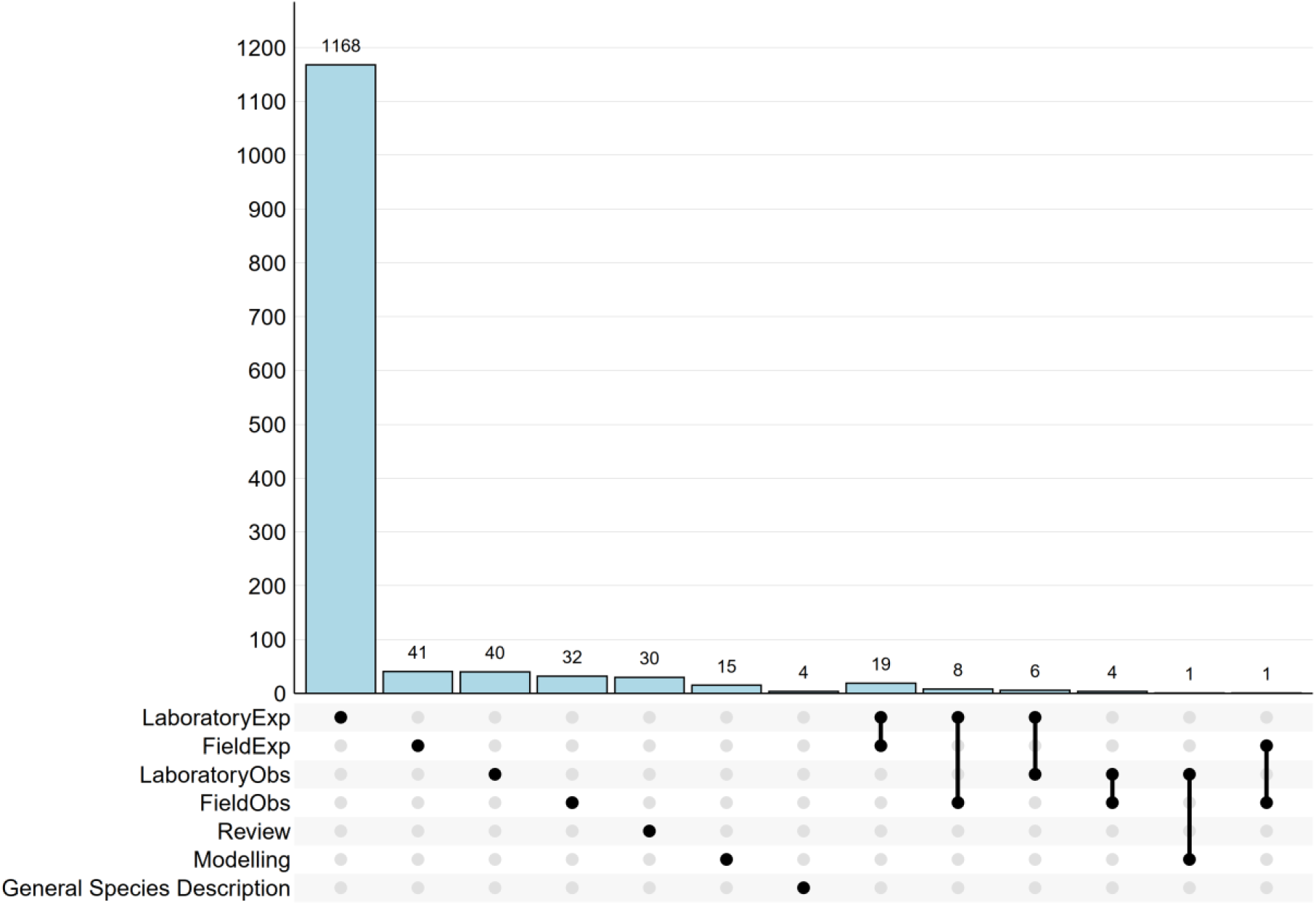
An UpSet plot of study design distribution in articles included after full-text screening. Each column corresponds to a unique intersection: filled, connected dots indicate the study design groups included; the bar above shows the number of articles with that study design.

**Figure 4.**
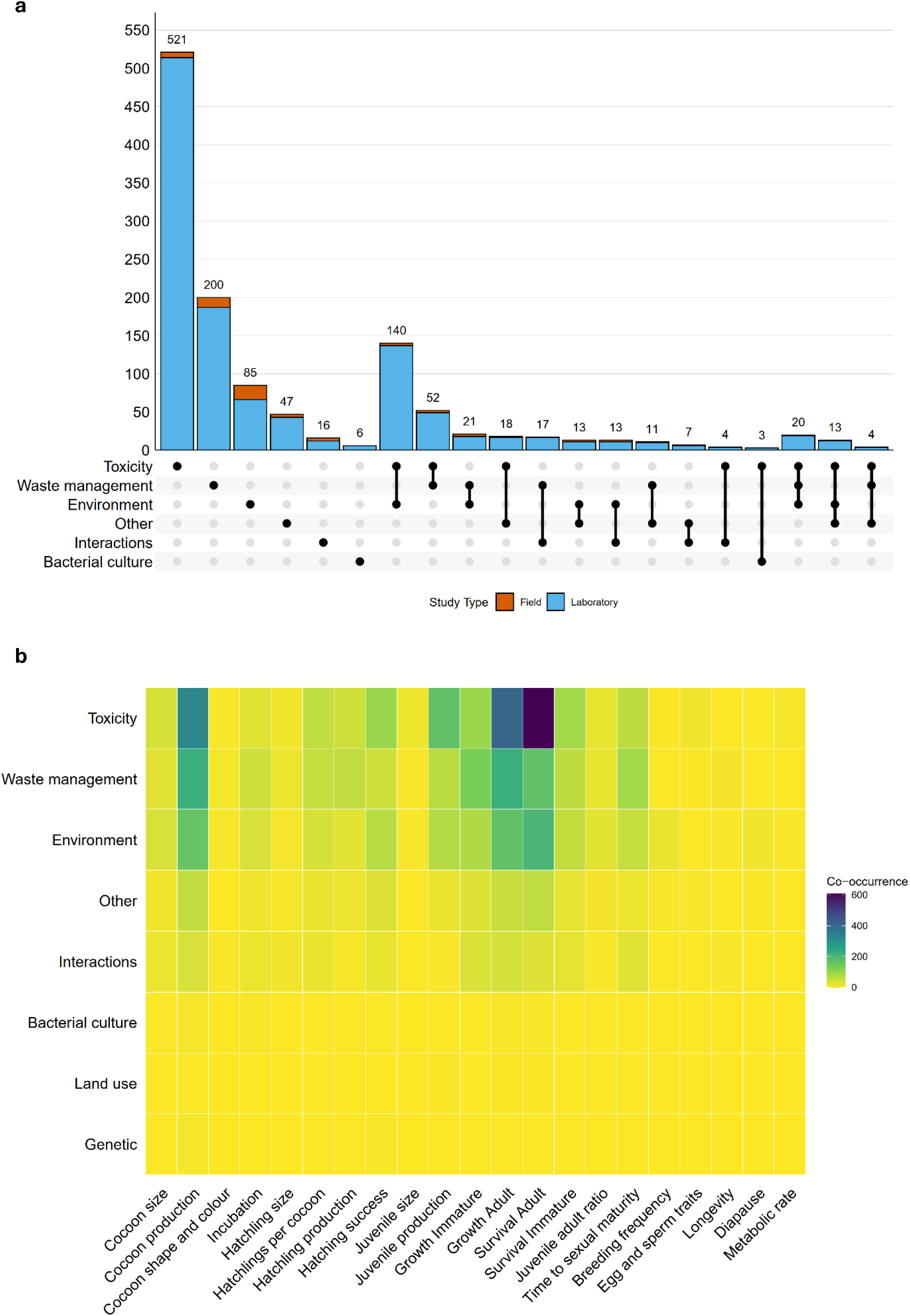
(a) UpSet plot of treatment group distribution; (b) heatmap of co-occurrence of treatment groups and life-history traits reported as response variables in articles included after full-text screening.

**Figure 5.**
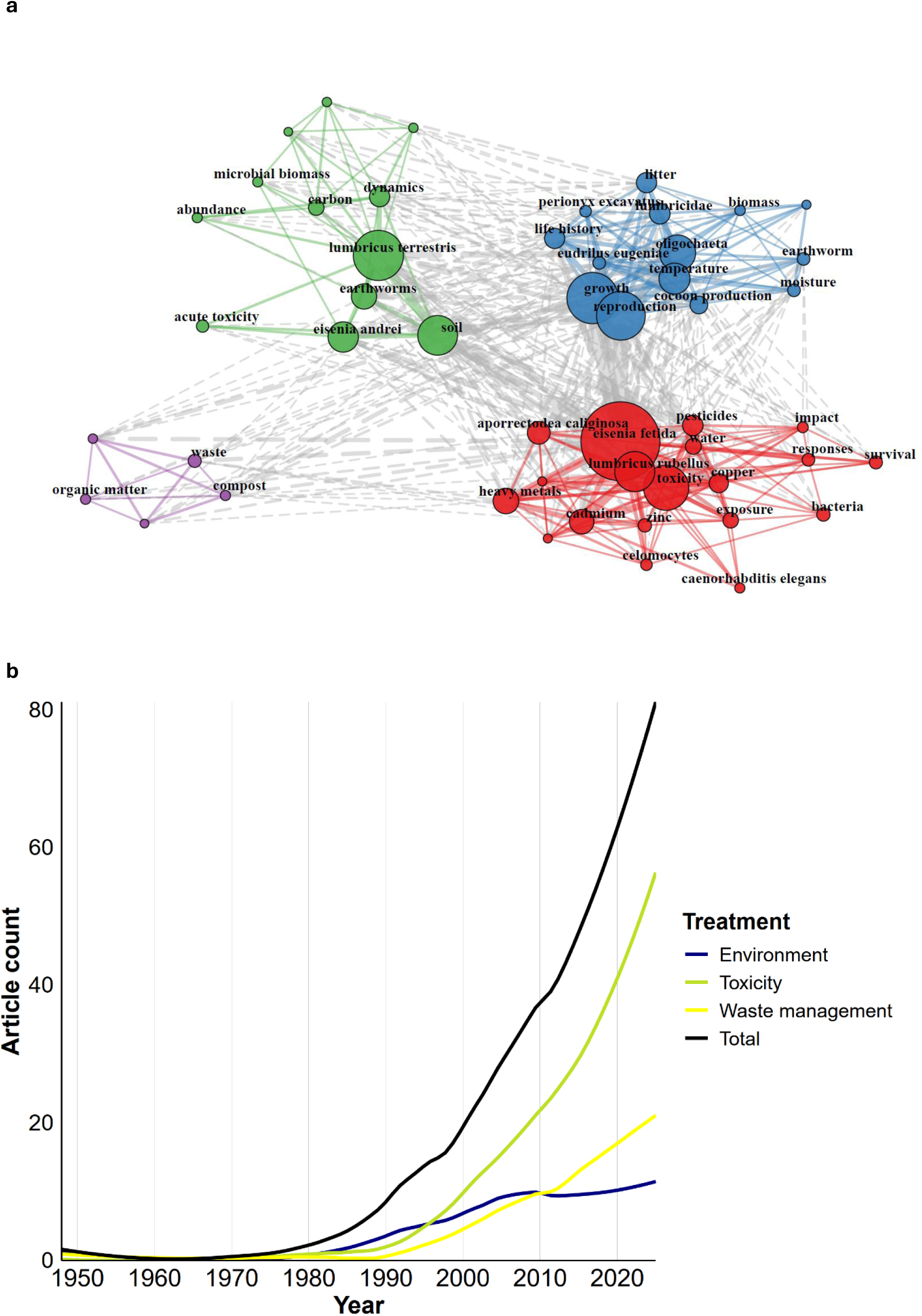
(a) Keyword co-occurrence map depicting clusters of keywords associated with articles included in full-text screening; (b) distribution of publications included in the full-text screening by year (LOESS smoothing with a 0.5 span applied to the article count).

**Figure 6.**
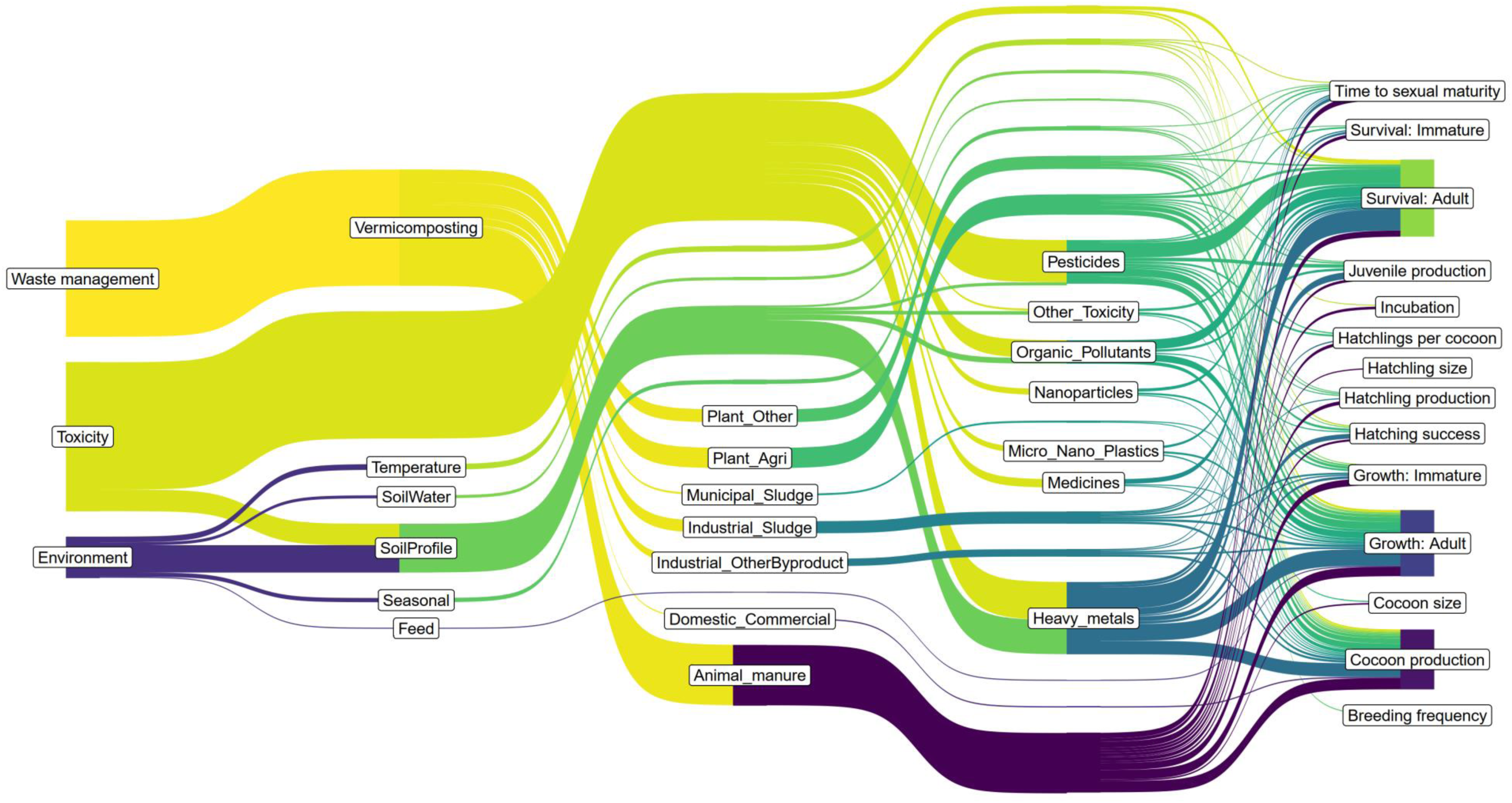
Sankey plot illustrating the connections between the main treatment groups, their subtypes and life-history trait outcomes. Only paths with more than 15 connections are displayed.

In total, studies on LH trait variation were carried out in 84 countries. India had the highest number of studies (*n* = 275), followed by China (*n* = 128) and United States (*n* = 94) (Fig 7; SM: Table S5). Asia, Europe and North America were the continents with the highest number of studies: 492, 457 and 170, respectively (SM: Table S6). This results in a bias of research effort towards the Northern Hemisphere as fewer studies have been carried out in South America (*n* = 89) and Oceania (*n* = 34). However, there were 106 LH studies from Africa, predominantly from South Africa. When broken down by study species, countries such as China, Spain and Brazil had a high number of articles, but these tended to focus exclusively on *Eisenia* spp. By contrast, as indicated in Fig. 7, LH studies carried out in France, Denmark, Finland and Egypt had a higher diversity of species studied.

**Figure 7.**
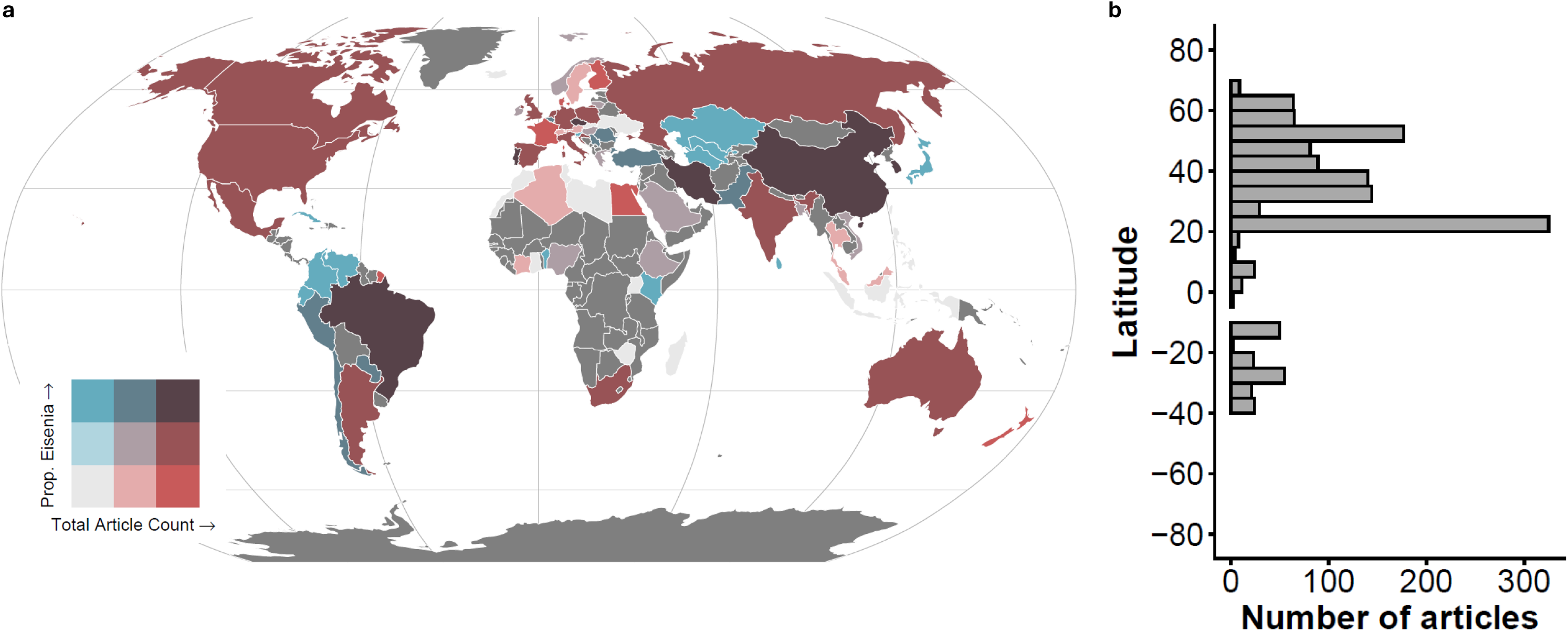
(a) geographical distribution of study locations, with total article counts on x axis and the proportion of articles exclusively using *Eisenia fetida (E. foetida*) *and/or E. andrei* as study systems on y axis; (b) histogram of article counts by latitudinal band.

## IV. DISCUSSION

Our systematic evidence map shows that research on earthworm life-history variation is unevenly distributed across species, traits, study designs and regions. The literature is dominated by studies on *E*. *fetida* and *E. andrei* and other introduced European species rather than native fauna, reflecting the widespread use of these taxa in laboratory-based research across the globe. The treatment groups identified in the experimental studies reviewed here generally map onto the framework of global change (GC) categories (Murphy & Romanuk, 2014; Phillips *et al*., 2024). The effect of pollution (‘chemical and physical toxicity’) and nutrient enrichment (‘waste management’) are particularly well studied, which highlights the importance of LH traits in understanding biological responses to these factors. By contrast, the direct impacts of invasive species (‘interactions’), climate change (‘environment’), land-use intensification (‘land use’) and habitat fragmentation on LH trait variation remain less understood. Because most studies were conducted under controlled conditions and addressed applied questions in the fields of ecotoxicology or organic waste management, they offer limited insight into how LH traits vary across natural environments. The geographical distribution in research effort also reflects hubs of activity in these fields, with studies concentrated in India and China. Despite the high number of articles from these countries, a large proportion of these articles focus exclusively on lab-based research in *Eisenia* model species.

### (1) Dominance of toxicity and waste management studies in LH research

As expected, our study highlights the outsized role of a few cosmopolitan species in LH research, particularly *E*. *fetida* and *E. andrei*, model organisms with a long history of use in ecotoxicology research (OECD, 1984, 2016; ISO, 2008). Species with high reproductive output and short life cycle such as these are commonly used in ecotoxicological testing because these LH traits make them easy to rear in the laboratory and enable rapid data collection (Yasmin & D’Souza, 2007; Vasseur & Bonnard, 2014). These same traits also contribute to their cosmopolitan status as they facilitate rapid population growth across diverse habitats. Meta-analyses indicate that *E. fetida* and *E. andrei* are generally less sensitive to heavy metals (Nahmani, Hodson & Black, 2007) and pesticides (Daam *et al*., 2011; Pelosi, Joimel & Makowski, 2013) compared to other earthworm species and other groups of soil fauna, but they show greater overall sensitivity to organic pollutants (Chao *et al*., 2022). In addition, *E*. *fetida* and *E. andrei* are typically found in compost or manure and rarely in mineral soils, which are most affected by pollutants (Lowe & Butt, 2007). Earthworms are commonly divided into epigeic, endogeic, and anecic groups based on differences in morphology, feeding strategy, and burrowing behaviour, corresponding broadly to surface-, shallow, and vertical-burrowing modes (Bouché, 1977). Because they occupy different parts of the soil profile, these groups differ in their effect on ecosystem processes (Blouin *et al*., 2013; Huang, González & Zou, 2020). Their abundance also varies with climate zones, for example, endogeic species are more abundant in the tropics while epigeic and anecic species dominate in boreal and temperate regions (Lavelle, 1983). Comprehensive ecotoxicological testing should include species from all three ecological categories and reflect the range of climates and soils they inhabit. Our current approaches, nevertheless, rely disproportionately on epigeic *Eisenia* species and data for anecic earthworms come mainly from temperate taxa such as *L*. *terrestris* (Medina-Sauza *et al*., 2019). The recent ISO standard 11268-2 (2023) for the assessment of pollutant effects on earthworms proposed more ecologically relevant species to be tested: epigeic *B. rubidus* for boreal, taiga and northern ecozone regions, and endogeic *A. caliginosa* as representative of temperate regions and agricultural soils. However, these are cosmopolitan species, and there remains an urgent need to test a wider range of species to understand what effects toxic chemicals and wastes might have on native earthworms with limited distribution (e.g., (Foerster, 2006; Gowri & Thangaraj, 2020; Zhang *et al*., 2022). The taxonomic bias towards a few model species identified in our study could also apply to other groups of soil fauna routinely used for toxicological testing. For example, *Folsomia candida* (collembolans), *Enchytraeus crypticus* (enchytraeids), and *Porcellio scaber* (isopods) (van Gestel, 2012) might have received disproportionately more research effort compared to other species in these clades. We join other recent calls to expand the taxonomic scope of ecotoxicity studies (e.g., Hecker, 2018;

Rosner *et al*., 2024; Roani *et al*., 2026), specifically to non-model organisms that can best approximate biological systems and evaluate long-term effects and ecosystem-wide impacts of environmental pollution. Reproductive output is an important endpoint used in assessing the sublethal effects of chemicals on earthworms (OECD, 2016) as reduction in fertility or fecundity can cause decrease in earthworm populations after several generations (Brunninger, Viswanathan & Beese, 1994). Indeed, our study reflects that importance, as reproductive output is widely reported, most commonly as cocoon production, hatching success and juvenile production. Reproductive endpoints measured in chronic tests are often markedly more sensitive to toxic substances than mortality endpoints from acute tests (e.g., 2–3 times more for Ni in *Eisenia veneta* (Scott-Fordsmand, Weeks & Hopkin, 1998)). In another meta-analysis, authors found that for 95% of reviewed cases for *E. fetida*, reproduction was more than twice as sensitive to pesticides as mortality, making it the more protective endpoint for risk assessment (Frampton *et al*., 2006). As a result, acute adult mortality test is no longer mandatory for pre-market regulatory testing in many countries (though it is still required in Brazil (IBAMA Normative Ordinance No. 84, 1996) and US (EPA Pesticide Registration Manual, 2026)), with chronic reproduction tests normally being a standard requirement in regulatory dossiers for pesticide registration (ISO standard 11268-2 (2023)). Although fecundity estimates are widely available for the species examined here, the dominance of cosmopolitan species would make it difficult to meaningfully compare reproductive trait variation between native and non-native species in the US and Brazil. Such comparisons would be important because invasive species are expected to exhibit higher reproductive outputs than native species. Longer-term studies of earthworm reproductive mode and phenology are also needed to establish breeding frequency, another trait likely to contribute to invasion success, as current research is largely constrained to established toxicological endpoints.

The majority of studies on survival and growth do not provide estimates for immature stages (i.e., hatchlings, juveniles, subadults) indicating that they are still under-researched compared to adults. Juvenile earthworms have been shown to be more sensitive to toxic substances than adults, suggesting that risk assessments using only adult specimens may underestimate harmful effects on earthworm populations (Van Gestel & Weeks, 2004; Pelosi *et al*., 2014). Moreover, this high sensitivity of immature stages is particularly concerning when it comes to species establishment, e.g., for *L*. *terrestris*, juvenile survival is a key driver of earthworm population dynamics (Pelosi *et al*., 2008). In contrast to other animal clades where egg size is a commonly used proxy for reproductive investment (Smith & Fretwell, 1974; Einum & Fleming, 2002), research on early earthworm development, i.e., eggs and sperm, has been the focus of relatively few studies (*n* = 25). In Bart *et al*. (2019), cocoon weight of *A. caliginosa* did not appear to be affected by pesticide exposure, suggesting that earthworms may adjust the number of cocoons produced rather than the amount of vitellus (i.e., yolk) allocated to each one. But this result cannot be generalised to all situations, as it remains to be confirmed for other substances and species. The limited attention given to immature stages reflects a broader issue in soil fauna research, where juveniles and adults of the same species are often treated as equivalent despite instances of developmental shifts in their ecological roles, as occurs, for instance, with predatory insect larvae shifting to phytophagous adults (Gongalsky, 2021). In earthworms, different developmental stages have been associated with variation in metabolic demands (Phillipson & Bolton, 1976; Uvarov & Scheu, 2004) and dietary preferences (Briones, 2001). Juvenile earthworms, however, may not occupy the same niches as adults, since they have been documented to feed near the soil surface irrespective of ecological group (Gerard, 1967; Rundgren, 1975). For instance, Pelosi *et al*., (2008) in their population dynamics model considered that juveniles of *L. terrestris* do not migrate vertically. They explained that these individuals are generally found close to the food source, within 8 cm of the soil surface, and that they do not yet have a sufficiently developed muscular system that Bouché (1977) called ‘digging muscles’. As juveniles can constitute > 50% of earthworm populations in annual cropping soils (e.g., Smith *et al*., 2008; Peigné *et al*., 2009; Pelosi *et al*., 2015; Briones & Schmidt, 2017; Bruz *et al*., 2023)), it is important to understand how their response to external factors might differ from adults.

### (2) Relevance for global change research

Anthropogenic disturbance is widely recognised as a key driver of invasion success, facilitating the establishment of non-native species across a range of taxa (e.g., Grarock *et al*., 2014; Salyer, Bennett & Buczkowski, 2014; Jauni, Gripenberg & Ramula, 2015). By altering native communities and creating novel ecological niches, land use change and climate warming may increase opportunities for non-native earthworms to establish and spread. In tropical habitats, slash-and-burn deforestation can substantially increase non-native earthworm populations, outweighing the effect of propagule pressure or food availability (He, 2020). In northern hardwood stands, liming of acidic soils has similarly been shown to increase the invasion risk of *L. terrestris* (Moore, Ouimet & Bohlen, 2013; Homan *et al*., 2016). At broader spatial scales, rising temperatures are expected to drive earthworm range expansions poleward and into higher elevations, with Arctic ecosystems becoming more vulnerable as permafrost thaws. European lumbricids such as *Aporrectodea* and *Lumbricus* spp. have already been recorded in the Arctic soils of North America, Fennoscandia and Russia (Hendrix *et al*., 2008; Blume-Werry *et al*., 2020; Mathieu *et al*., 2024). Given the growing importance of understanding and predicting the effects of multiple global change stressors acting in tandem (Rillig *et al*., 2019; Bowler *et al*., 2020), it is promising that our analysis identified a number of studies examining the effects of multiple treatment types on LH traits (e.g., interaction between temperature and heavy metal toxicity (Svendsen *et al*., 2007) or between plant presence and cadmium toxicity (Du *et al*., 2014); management of transgenic maize waste (Zhang *et al*., 2025)).

The exponential increase in research related to toxicity and waste management in the 1980’s and 1990’s, respectively, likely reflects the emergence of standardized testing frameworks and regulatory requirements for assessing effects on earthworms (OECD, 1984; ISO 11268-1, 1993). We do note that, in contrast to these two treatment types, the number of articles on environmental factors has not been subject to exponential growth over the last decades, even though research on biodiversity loss is dominated by studies on climate change impact (Mazor *et al*., 2018). Some groups of organisms, such as ectotherms, may be particularly at risk from climate change impacts including elevated temperatures and drought because they depend mainly on behavioural strategies rather than internal physiological regulation to avoid stressful environments (Kingsolver, Diamond & Buckley, 2013; Walsh *et al*., 2019). Earthworms are ectotherms that breathe through the skin, making them highly sensitive to environmental conditions (Edwards & Arancon, 2022). Their vulnerability is further increased by their small body size, as their internal temperature changes more quickly in response to ambient conditions than that of larger animals (Stevenson, 1985). Experimental studies reviewed here show that the interaction between temperature and soil moisture exerts a strong influence on LH traits (e.g., Reinecke & Kriel, 1981; Reinecke, Viljoen & Saayman, 1992; Holmstrup, 2001). However, we are still lacking a large-scale synthesis of environmental impacts on earthworm traits.

We found few field studies exploring the impact of GC stressors on LH traits. The lack of studies investigating the effect of land-use intensification, particularly tillage, is especially concerning given the well-documented negative impacts of this practice on earthworm communities (Briones & Schmidt, 2017). Field studies provide insights into LH trait variation under more realistic ecological conditions but remain scarce due to methodological constraints. First, measuring LH traits *in situ* requires repeated detection of individuals, which is challenging for earthworms once they are buried in soil. Although mark–recapture methods are widely used to monitor animal populations, they are difficult to apply to earthworms (Mathieu, Caro & Dupont, 2018). Moreover, surveying the age structure of field populations to obtain per-species estimates of juvenile:adult ratio is challenging due to difficulties with taxonomic identification of juveniles, as well as the considerable sampling effort needed to assess population trends over time. GC impact might also differ by life stage, e.g., juveniles are particularly sensitive to tillage (Ulrich *et al*., 2010; Briones & Schmidt, 2017). DNA barcoding can help address these issues by enabling more reliable species identification during early life stages and at scale (Richard *et al*., 2010; Decaëns *et al*., 2013; Eydoux *et al*., 2026), but remains rather costly in some countries, making its widespread use still limited. A test guideline (OECD, 2026) has been developed for assessing short- and long-term effects of chemicals on earthworm communities in soils under field conditions. However, this guideline was primarily developed for prospective regulatory risk assessment by manufacturers prior to market authorisation and, owing to its complexity, cost and logistical constraints, has so far seen limited routine application outside regulatory contexts.

Finally, relatively few studies have examined interactions between species, and even fewer have estimated the direct impacts of invasive species on native species (e.g., Ortiz-Ceballos, 2005; Winsome *et al*., 2006). Because species are most often studied in isolation, such approaches may fail to capture the complexity of biotic interactions in nature. For example (Winsome *et al*., 2006) compared the native Californian species *Argilophilus marmoratus* with non-native *Aporrectodea trapezoides* in field and laboratory experiments by varying grassland productivity and competitor density. *A. trapezoides* reproduced much faster, with many individuals becoming reproductive within weeks of emergence, giving it a potential competitive advantage over the native species; however, the native species had a negative effect on the invasive species in a low productivity setting. In a study of invasive lumbricids in Western Siberia (Golovanova *et al*., 2023), the juvenile and adult numbers of native species *Eisenia nordenskioldi* was higher in the presence of the invasive *A*. *caliginosa* compared to a monoculture, which highlights that the impact of non-native species is not always negative. To understand the broader consequences of invasive earthworms for ecosystem functioning, future studies should examine their interactions with the wider soil community, including micro-, meso-, and other macrofauna, microbes, fungi and plants. In temperate latitudes, invasive *Amynthas* spp. have been shown to affect both lower and higher trophic levels through their effects on collembolans and their role as novel prey for centipedes (Gao, Taylor & Callaham, 2017) and garter snakes (Crone *et al*., 2022). Expanding this research to tropical and subtropical regions would be important, as these areas host a different set of invasive species and a large number of native species.

## V. CONCLUSIONS

(1) Our study revealed a large body of research on earthworm LH traits; however, this research is highly biased. More studies on native species are urgently needed, particularly in tropical habitats, to better understand the impacts of environmental disturbances on earthworm communities. Greater attention should also be given to tropical and subtropical introduced species, considering the widespread endemicity and land-conversion processes occurring throughout this region.
(2) To improve ecological relevance, toxicity testing should use a broader range of model earthworm species that reflects both taxonomic diversity and variation in ecological groups, under different soil and climatic conditions.
(3) Due to the importance of juvenile stages in sustaining earthworm populations, greater attention should be given to immature stages to have a deeper understanding of population dynamics.
(4) More research on environmental impacts, e.g., temperature and soil moisture, on earthworm life histories in natural environments is needed, especially on native species. This is especially important given the widespread impacts of global change.
(5) To better capture the effect of biotic interactions on earthworms in nature, future studies should assess the direct effects of inter- and intra-specific interactions.

## VI. ACKNOWLEDGEMENTS

KV, HRPP – University of Helsinki. KN, DR, SG, HRPP – Research Council of Finland Grant (Number 362759) to HRPP. EKC acknowledges the support of the Natural Sciences and Engineering Research Council of Canada and the Canada Research Chairs program (reference numbers: RGPIN-2019-05758 and CRC-2024-00223).

## VIII. DATA AVAILABILITY STATEMENT

Datasets and code used in this work will be available in Zenodo (doi to be provided) and GitHub (link to be provided).

## Notes

### Competing Interest Statement

The authors have declared no competing interest.

## REFERENCES

Ahlmann-Eltze, C. (2025) ggupset: Combination Matrix Axis for ‘ggplot2’ to Create ‘UpSet’ Plots. https://cran.r-project.org/web/packages/ggupset/index.html [accessed 18 May 2026].

Allen, W.L., Street, S.E. & Capellini, I. (2017) Fast life history traits promote invasion success in amphibians and reptiles. Ecology Letters 20, 222–230.

Anthony, M.A., Bender, S.F. & van der Heijden, M.G.A. (2023) Enumerating soil biodiversity. Proceedings of the National Academy of Sciences 120, e2304663120.

Aria, M. & Cuccurullo, C. (2017) *bibliometrix*: An R-tool for comprehensive science mapping analysis. Journal of Informetrics 11, 959–975.

Bart, S., Barraud, A., Amosse, J., Pery, A.R.R., Mougin, C. & Pelosi, C. (2019) Effects of two common fungicides on the reproduction of *Aporrectodea caliginosa* in natural soil. Ecotoxicology and Environmental Safety 181, 518–524.

Baumann, T.T., Frelich, L.E., Van Riper, L.C. & Yoo, K. (2024) Anthropogenic transport mechanisms of invasive European earthworms: a review. Biological Invasions 26, 3563–3586.

Bello, F. de, Lavorel, S., Hallett, L.M., Valencia, E., Garnier, E., Roscher, C., Conti, L., Galland, T., Goberna, M., Májeková, M., Montesinos-Navarro, A., Pausas, J.G., Verdú, M., E-Vojtkó, A., Götzenberger, L., et al. (2021) Functional trait effects on ecosystem stability: assembling the jigsaw puzzle. Trends in Ecology & Evolution 36, 822–836.

Blackburn, T.M., Essl, F., Evans, T., Hulme, P.E., Jeschke, J.M., Kühn, I., Kumschick, S., Marková, Z., Mrugała, A., Nentwig, W., Pergl, J., Pyšek, P., Rabitsch, W., Ricciardi, A., Richardson, D.M., et al. (2014) A unified classification of alien species based on the magnitude of their environmental impacts. PLOS Biology 12, e1001850.

Blackburn, T.M., Lockwood, J.L. & Cassey, P. (2015) The influence of numbers on invasion success. Molecular Ecology 24, 1942–1953.

Blakemore, R.J. (2009). Cosmopolitan Earthworms—A Global and Historical Perspective. In Annelids in Modern Biology, D.H. Shain (Ed.), John Wiley & Sons.

Blouin, M., Hodson, M.E., Delgado, E.A., Baker, G., Brussaard, L., Butt, K.R., Dai, J., Dendooven, L., Peres, G., Tondoh, J.E., Cluzeau, D. & Brun, J.-J. (2013) A review of earthworm impact on soil function and ecosystem services. European Journal of Soil Science 64, 161–182.

Blume-Werry, G., Krab, E.J., Olofsson, J., Sundqvist, M.K., Väisänen, M. & Klaminder, J. (2020) Invasive earthworms unlock arctic plant nitrogen limitation. Nature Communications 11, 1766.

Bouché, M.B. (1977) Strategies lombriciennes. Ecological Bulletins 25, 122–132.

Bowler, D.E., Bjorkman, A.D., Dornelas, M., Myers-Smith, I.H., Navarro, L.M., Niamir, A., Supp, S.R., Waldock, C., Winter, M., Vellend, M., Blowes, S.A., Böhning-Gaese, K., Bruelheide, H., Elahi, R., Antão, L.H., et al. (2020) Mapping human pressures on biodiversity across the planet uncovers anthropogenic threat complexes. People and Nature 2, 380–394.

Bradshaw, C.J.A., Hulme, P.E., Hudgins, E.J., Leung, B., Kourantidou, M., Courtois, P., Turbelin, A.J., McDermott, S.M., Lee, K., Ahmed, D.A., Latombe, G., Bang, A., Bodey, T.W., Haubrock, P.J., Saltré, F., et al. (2024) Damage costs from invasive species exceed management expenditure in nations experiencing lower economic activity. Ecological Economics 220, 108166.

Briones, M.J.I. & Schmidt, O. (2017) Conventional tillage decreases the abundance and biomass of earthworms and alters their community structure in a global meta-analysis. Global Change Biology 23, 4396–4419.

Briones, R.S., MJI Bol (2001) Spatio-temporal variation of stable isotope ratios in earthworms under grassland and maize cropping systems. Soil Biology & Biochemistry 33, 1673–1682.

Brown, G., Feller, C., Blanchart, E., Deleporte, P. & Chernyanskii, S. (2003) With Darwin, earthworms turn intelligent and become human friends. Pedobiologia 47, 924–933.

Brunninger, B., Viswanathan, R. & Beese, F. (1994) Terbuthylazine and carbofuran effects on growth and reproduction within 3 generations of *Eisenia andrei* (Oligochaeta). Biology and Fertility of Soils 18, 83–88.

Bruz, L.D.S.M., Santos, A., Demetrio, W.C., Primo, D.D.F., Feliciano, L.P., Fernandes, C.H., Bartz, M.L.C., Bernardi, A.C.D.C., Pezzopane, J.R.M. & Brown, G.G. (2023) Earthworms in various agricultural and forest ecosystems in São Carlos—SP, Brazil. Zootaxa 5255, 324–335.

Cameron, E., Martins, I.S., Lavelle, P., Mathieu, J., Tedersoo, L., Bahram, M., Gottschall, F., Guerra, C., Hines, J., Patoine, G., Siebert, J., Winter, M., Cesarz, S., Ferlian, O., Kreft, H., et al. (2019) Global mismatches in aboveground and belowground biodiversity. Conservation Biology 33, 1187–1192.

Cameron, E.K. & Bayne, E.M. (2015) Spatial patterns and spread of exotic earthworms at local scales. Canadian Journal of Zoology 93, 721–726.

Cameron, E.K., Bayne, E.M. & Clapperton, M.J. (2007) Human-facilitated invasion of exotic earthworms into northern boreal forests. Écoscience 14, 482–490.

Cameron, E.K., Martins, I.S., Lavelle, P., Mathieu, J., Tedersoo, L., Gottschall, F., Guerra, C.A., Hines, J., Patoine, G., Siebert, J., Winter, M., Cesarz, S., Delgado-Baquerizo, M., Ferlian, O., Fierer, N., et al. (2018) Global gaps in soil biodiversity data. Nature Ecology & Evolution 2, 1042–1043.

Capellini, I., Baker, J., Allen, W.L., Street, S.E. & Venditti, C. (2015) The role of life history traits in mammalian invasion success. Ecology Letters 18, 1099–1107.

Chang, C.-H., Bartz, M.L.C., Brown, G., Callaham, M.A., Jr., Cameron, E.K., Davalos, A., Dobson, A., Gorres, J.H., Herrick, B.M., Ikeda, H., James, S.W., Johnston, M.R., McCay, T.S., McHugh, D., Minamiya, Y., et al. (2021) The second wave of earthworm invasions in North America: biology, environmental impacts, management and control of invasive jumping worms. Biological Invasions 23, 3291–3322.

Chao, H., Sun, M., Wu, Y., Xia, R., Yuan, S. & Hu, F. (2022) Quantitative relationship between earthworms’ sensitivity to organic pollutants and the contaminants’ degradation in soil: A meta-analysis. Journal of Hazardous Materials 429, 128286.

Chauvel, A., Grimaldi, M., Barros, E., Blanchart, E., Desjardins, T., Sarrazin, M. & Lavelle, P. (1999) Pasture damage by an Amazonian earthworm. Nature 398, 32–33.

Chown, S.L. & McGeoch, M.A. (2023) Functional trait variation along animal invasion pathways. Annual Review of Ecology, Evolution, and Systematics 54, 151–170.

Corporation for Digital Scholarship (2023) Zotero. https://www.zotero.org/.

Costello, D.M., Tiegs, S.D. & Lamberti, G.A. (2011) Do non-native earthworms in Southeast Alaska use streams as invasional corridors in watersheds harvested for timber? Biological Invasions 13, 177–187.

Craven, D., Thakur, M.P., Cameron, E.K., Frelich, L.E., Beauséjour, R., Blair, R.B., Blossey, B., Burtis, J., Choi, A., Dávalos, A., Fahey, T.J., Fisichelli, N.A., Gibson, K., Handa, I.T., Hopfensperger, K., et al. (2017) The unseen invaders: introduced earthworms as drivers of change in plant communities in North American forests (a meta-analysis). Global Change Biology 23, 1065–1074.

Crone, E.R., Sauer, E.L., Herrick, B.M., Drake, D. & Preston, D.L. (2022) Effects of invasive jumping worms (*Amynthas* spp.) on microhabitat and trophic interactions of native herpetofauna. Biological Invasions 24, 2499–2512.

Cuthbert, R.N., Bodey, T.W., Briski, E., Capellini, I., Dick, J.T.A., Kourantidou, M., Ricciardi, A. & Pincheira-Donoso, D. (2025) Harnessing traits to predict economic impacts from biological invasions. Trends in Ecology & Evolution 40, 639–650.

Cuthbert, R.N., Diagne, C., Hudgins, E.J., Turbelin, A., Ahmed, D.A., Albert, C., Bodey, T.W., Briski, E., Essl, F., Haubrock, P.J., Gozlan, R.E., Kirichenko, N., Kourantidou, M., Kramer, A.M. & Courchamp, F. (2022) Biological invasion costs reveal insufficient proactive management worldwide. Science of The Total Environment 819, 153404.

Daam, M.A., Leitao, S., Cerejeira, M.J. & Paulo Sousa, J. (2011) Comparing the sensitivity of soil invertebrates to pesticides with that of *Eisenia fetida*. Chemosphere 85, 1040–1047.

Decaëns, T. (2010) Macroecological patterns in soil communities. Global Ecology and Biogeography 19, 287–302.

Decaëns, T., Jiménez, J.J., Gioia, C., Measey, G.J. & Lavelle, P. (2006) The values of soil animals for conservation biology. European Journal of Soil Biology 42, S23–S38.

Decaëns, T., Porco, D., Rougerie, R., Brown, G.G. & James, S.W. (2013) Potential of DNA barcoding for earthworm research in taxonomy and ecology. Applied Soil Ecology 65, 35–42.

Demetrio, W., Brown, G., Pupin, B., Novo, R., Dudas, R., Baretta, D., Römbke, J., Bartz, M. & Borma, L. (2023) Are exotic earthworms threatening soil biodiversity in the Brazilian Atlantic Forest? Applied Soil Ecology 182, 104693.

Dempsey, M.A., Fisk, M.C., Yavitt, J.B., Fahey, T.J. & Balser, T.C. (2013) Exotic earthworms alter soil microbial community composition and function. Soil Biology & Biochemistry 67, 263–270.

Du, Y.-L., He, M.-M., Xu, M., Yan, Z.-G., Zhou, Y.-Y., Guo, G.-L., Nie, J., Wang, L.-Q., Hou, H. & Li, F.-S. (2014) Interactive effects between earthworms and maize plants on the accumulation and toxicity of soil cadmium. Soil Biology & Biochemistry 72, 193–202.

Edwards, C.A. & Arancon, N.Q. (2022) Earthworm Life Histories and Biology. In Biology and Ecology of Earthworms pp. 81–108. Springer US, New York, NY.

Einum, S. & Fleming, I.A. (2002) Does within-population variation in fish egg size reflect maternal influences on optimal values? The American Naturalist 160, 756–765.

Eisenhauer, N., Partsch, S., Parkinson, D. & Scheu, S. (2007) Invasion of a deciduous forest by earthworms: Changes in soil chemistry, microflora, microarthropods and vegetation. Soil Biology and Biochemistry 39, 1099–1110.

EPA (2026) Pesticide Registration Manual. Office of Pesticide Programs

Eydoux, L., Hedde, M., Valentini, A., Decaëns, T., Lafranchi, Q., Lucas, A., Jay-Robert, P. & Vergnes, A. (2026) Beneath the surface: exploring earthworm assemblages in recently unsealed urban schoolyards using eDNA metabarcoding. Applied Soil Ecology 223, 107113.

Foerster, M.F., Bernhard Garcia (2006) Effects of carbendazim and lambda-cyhalothrin on soil invertebrates and leaf litter decomposition in semi-field and field tests under tropical conditions (Amazonia, Brazil). European Journal of Soil Biology 42, S171–S179.

Foo, Y.Z., O’Dea, R.E., Koricheva, J., Nakagawa, S. & Lagisz, M. (2021) A practical guide to question formation, systematic searching and study screening for literature reviews in ecology and evolution. Methods in Ecology and Evolution 12, 1705–1720.

Frampton, G.K., Jaensch, S., Scott-Fordsmand, J.J., Roembke, J. & Van den Brink, P.J. (2006) Effects of pesticides on soil invertebrates in laboratory studies: A review and analysis using species sensitivity distributions. Environmental Toxicology and Chemistry 25, 2480–2489.

Froese, R. & Pauly, D. (2025) FishBase. https://www.fishbase.se/ [accessed 20 August 2025].

Gao, M., Taylor, M.K. & Callaham, Jr., Mac A. (2017) Trophic dynamics in a simple experimental ecosystem: Interactions among centipedes, Collembola and introduced earthworms. Soil Biology and Biochemistry 115, 66–72.

Gerard, B. (1967) Factors affecting earthworms in pastures. Journal of Animal Ecology 36, 235–252.

van Gestel, C.A.M. (2012) Soil ecotoxicology: state of the art and future directions. ZooKeys 176, 275–296.

Golovanova, E.V., Kniazev, S.Y., Karaban, K., Babiy, K.A. & Shekhovtsov, S.V. (2023) First short-term study of the relationship between native and invasive earthworms in the zone of soil freezing in Western Siberia-experiments in mesocosms. Diversity-Basel 15, 248.

Gongalsky, K.B. (2021) Soil macrofauna: Study problems and perspectives. Soil Biology and Biochemistry 159, 108281.

Görres, J.H., Bellitürk, K. & Melnichuk, R.D.S. (2016) Temperature and moisture variables affecting the earthworms of genus *Amynthas* Kinberg, 1867 (Oligachaeta: Megascolecidae) in a hardwood forest in the Champlain Valley, Vermont, USA. Applied Soil Ecology 104, 111–115.

Gowri, S. & Thangaraj, R. (2020) Studies on the toxic effects of agrochemical pesticide (Monocrotophos) on physiological and reproductive behavior of indigenous and exotic earthworm species. International Journal of Environmental Health Research 30, 212–225.

Grames, E.M., Stillman, A.N., Tingley, M.W. & Elphick, C.S. (2019) An automated approach to identifying search terms for systematic reviews using keyword co-occurrence networks. Methods in Ecology and Evolution 10, 1645–1654.

Grarock, K., Tidemann, C.R., Wood, J.T. & Lindenmayer, D.B. (2014) Are invasive species drivers of native species decline or passengers of habitat modification? A case study of the impact of the common myna (*Acridotheres tristis*) on Australian bird species. Austral Ecology 39, 106–114.

Green, S.J., Brookson, C.B., Hardy, N.A. & Crowder, L.B. (2022) Trait-based approaches to global change ecology: moving from description to prediction. Proceedings of the Royal Society B: Biological Sciences 289, 20220071.

He, S.W., Xinxing Liu (2020) Disturbance intensity overwhelms propagule pressure and litter resource in controlling the success of *Pontoscolex corethrurus* invasion in the tropics. Biological Invasions 22, 1705–1721.

Hecker, M. (2018) Non-model Species in Ecological Risk Assessment. In A Systems Biology Approach to Advancing Adverse Outcome Pathways for Risk Assessment (eds N. Garcia-Reyero & C.A. Murphy), pp. 107–132. Springer International Publishing, Cham.

Hendrix, P.F. & Bohlen, P.J. (2002) Exotic earthworm invasions in North America: ecological and policy implications. BioScience 52, 801–811.

Hendrix, P.F., Jr, M.A.C., Drake, J.M., Huang, C.-Y., James, S.W., Snyder, B.A. & Zhang, W. (2008) Pandora’s box contained bait: The global problem of introduced earthworms*. Annual Review of Ecology, Evolution, and Systematics 39, 593–613.

Hodgins, K., Bock, D. & Rieseberg, L. (2018) Trait evolution in invasive species. Annual Plant Reviews Online 1, 459–496.

Holmstrup, M. (1995) Polyol accumulation in earthworm cocoons induced by dehydration. Comparative Biochemistry And Physiology A-Molecular & Integrative Physiology 111, 251–255.

Holmstrup, M. (2001) Sensitivity of life history parameters in the earthworm *Aporrectodea caliginosa* to small changes in soil water potential. Soil Biology & Biochemistry 33, 1217–1223.

Holmstrup, M. (2003) Overwintering adaptations in earthworms. Pedobiologia 47, 504–510.

Holmstrup, M. & Westh, P. (1994) Dehydration of earthworm cocoons exposed to cold - a novel cold-hardiness mechanism. Journal Of Comparative Physiology B-Biochemical Systemic And Environmental Physiology 164, 312–315.

Holmstrup & Westh (1995) Effects of dehydration on water relations and survival of Lumbricid earthworm egg capsules. Journal of Comparative Physiology B 165, 377–383.

Homan, C., Beier, C., McCay, T. & Lawrence, G. (2016) Application of lime (CaCO_3_) to promote forest recovery from severe acidification increases potential for earthworm invasion. Forest Ecology and Management 368, 39–44.

Huang, W., González, G. & Zou, X. (2020) Earthworm abundance and functional group diversity regulate plant litter decay and soil organic carbon level: A global meta-analysis. Applied Soil Ecology 150, 103473.

Hulme, P.E. (2017) Climate change and biological invasions: evidence, expectations, and response options. Biological Reviews 92, 1297–1313.

IBAMA. (1996, October 15). Portaria Normativa n° 84 [Normative Ordinance No. 84]. Diário Oficial da União.

ISO (1993) ISO 11268-1: Soil quality - Effects of pollutants on earthworms (*Eisenia fetida*)-Part 1: Determination of acute toxicity using artificial soil substrate. ISO Geneva, Switzerland.

ISO (2008) 17512-1: Soil quality-Avoidance test for determining the quality of soils and effects of chemicals on behaviour-Part 1: Test with earthworms (*Eisenia fetida* and *Eisenia andrei*). ISO Geneva, Switzerland.

ISO (2023) 11268-2: Soil quality - Effects of pollutants on earthworms Part 2: Determination of effects on reproduction of *Eisenia fetida/Eisenia andrei* and other earthworm species. ISO Geneva, Switzerland.

James, K.L., Randall, N.P. & Haddaway, N.R. (2016) A methodology for systematic mapping in environmental sciences. Environmental Evidence 5, 7.

Jauni, M., Gripenberg, S. & Ramula, S. (2015) Non-native plant species benefit from disturbance: a meta-analysis. Oikos 124, 122–129.

Joimel, S., Potapov, A., Pey, B., Bonfanti, J., Cortet, J., Almeida, T.D., Lonardo, S.D., Hackenberger, D.K., Krogh, P.H., Laskowski, R., Loureiro, S. & Hedde, M. (2024) Trait concepts, categories, and databases in soil invertebrate ecology – ordering the mess. Soil Organisms 96, 151–166.

Jones, K.E., Bielby, J., Cardillo, M., Fritz, S.A., O’Dell, J., Orme, C.D.L., Safi, K., Sechrest, W., Boakes, E.H., Carbone, C., Connolly, C., Cutts, M.J., Foster, J.K., Grenyer, R., Habib, M., et al. (2009) PanTHERIA: a species-level database of life history, ecology, and geography of extant and recently extinct mammals. Ecology 90, 2648–2648.

Kattge, J., Díaz, S., Lavorel, S., Prentice, I.C., Leadley, P., Bönisch, G., Garnier, E., Westoby, M., Reich, P.B., Wright, I.J., Cornelissen, J.H.C., Violle, C., Harrison, S.P., Van Bodegom, P.M., Reichstein, M., et al. (2011) TRY – a global database of plant traits. Global Change Biology 17, 2905–2935.

Kingsolver, J.G., Diamond, S.E. & Buckley, L.B. (2013) Heat stress and the fitness consequences of climate change for terrestrial ectotherms. Functional Ecology 27, 1415–1423.

Kumschick, S., Gaertner, M., Vilà, M., Essl, F., Jeschke, J.M., Pyšek, P., Ricciardi, A., Bacher, S., Blackburn, T.M., Dick, J.T.A., Evans, T., Hulme, P.E., Kühn, I., Mrugała, A., Pergl, J., et al. (2015) Ecological impacts of alien species: Quantification, scope, caveats, and recommendations. BioScience 65, 55–63.

Lavelle, P. (1983) The structure of earthworm communities. In Earthworm Ecology: From Darwin to Vermiculture (ed J.E. Satchell), pp. 449–466. Springer Netherlands, Dordrecht.

Ligthart, T. & Peek, G. (1997) Evolution of earthworm burrow systems after inoculation of lumbricid earthworms in a pasture in the Netherlands. Soil Biology & Biochemistry 29, 453–462.

Lockwood, J.L., Cassey, P. & Blackburn, T. (2005) The role of propagule pressure in explaining species invasions. Trends in Ecology & Evolution 20, 223–228.

Loss, S.R. & Blair, R.B. (2014) Earthworm invasions and the decline of clubmosses (*Lycopodium* spp.) that enhance nest survival rates of a ground-nesting songbird. Forest Ecology and Management 324, 64–71.

Lowe, C.N. & Butt, K.R. (2007) Earthworm culture, maintenance and species selection in chronic ecotoxicological studies: A critical review. European Journal of Soil Biology 43, S281–S288.

Lu, J.-Z., Pfingstl, T., Junker, R.R., Maraun, M., Erktan, A. & Scheu, S. (2025) Life history traits in microarthropods: Evidence for a soil animal economics spectrum. Geoderma 455, 117206.

Marinissen, J.C.Y. & van den Bosch, F. (1992) Colonization of new habitats by earthworms. Oecologia 91, 371–376.

Martin, L.J., Blossey, B. & Ellis, E. (2012) Mapping where ecologists work: biases in the global distribution of terrestrial ecological observations. Frontiers in Ecology and the Environment 10, 195–201.

Mathieu, J., Caro, G. & Dupont, L. (2018) Methods for studying earthworm dispersal. Applied Soil Ecology 123, 339–344.

Mathieu, J., Reynolds, J.W., Fragoso, C. & Hadly, E. (2024) Multiple invasion routes have led to the pervasive introduction of earthworms in North America. Nature Ecology & Evolution 8, 489–499.

Mazor, T., Doropoulos, C., Schwarzmueller, F., Gladish, D.W., Kumaran, N., Merkel, K., Di Marco, M. & Gagic, V. (2018) Global mismatch of policy and research on drivers of biodiversity loss. Nature Ecology & Evolution 2, 1071–1074.

McClain, C.R., Heim, N.A., Knope, M.L., Monarrez, P.M., Payne, J.L., Santos, I.T. & Webb, T.J. (2025) MOBS 1.0: A Database of Interspecific Variation in Marine Organismal Body Sizes. Global Ecology and Biogeography 34, e70062.

Medina-Sauza, R.M., Álvarez-Jiménez, M., Delhal, A., Reverchon, F., Blouin, M., Guerrero-Analco, J.A., Cerdán, C.R., Guevara, R., Villain, L. & Barois, I. (2019) Earthworms building up soil microbiota, a review. Frontiers in Environmental Science 7.

Moore, J.-D., Ouimet, R. & Bohlen, P.J. (2013) Effects of liming on survival and reproduction of two potentially invasive earthworm species in a northern forest Podzol. Soil Biology and Biochemistry 64, 174–180.

Murphy, G.E.P. & Romanuk, T.N. (2014) A meta-analysis of declines in local species richness from human disturbances. Ecology and Evolution 4, 91–103.

Nahmani, J., Hodson, M.E. & Black, S. (2007) Effects of metals on life cycle parameters of the earthworm *Eisenia fetida* exposed to field-contaminated, metal-polluted soils. Environmental Pollution 149, 44–58.

Neyret, M., Le Provost, G., Boesing, A.L., Schneider, F.D., Baulechner, D., Bergmann, J., de Vries, F.T., Fiore-Donno, A.M., Geisen, S., Goldmann, K., Merges, A., Saifutdinov, R.A., Simons, N.K., Tobias, J.A., Zaitsev, A.S., et al. (2024) A slow-fast trait continuum at the whole community level in relation to land-use intensification. Nature Communications 15, 1251.

O’Dea, R.E., Lagisz, M., Jennions, M.D., Koricheva, J., Noble, D.W.A., Parker, T.H., Gurevitch, J., Page, M.J., Stewart, G., Moher, D. & Nakagawa, S. (2021) Preferred reporting items for systematic reviews and meta-analyses in ecology and evolutionary biology: a PRISMA extension. Biological Reviews 96, 1695–1722.

OECD (1984) Test No. 207: Earthworm, acute toxicity tests. OECD Guidelines for the Testing of Chemicals. OECD Publishing.

OECD (2016) Test No. 222: Earthworm reproduction test (*Eisenia fetida/Eisenia andrei*). OECD Guidelines for the Testing of Chemicals, Section 2. OECD Publishing.

OECD (2026) Test No. 256: Determination of effects on earthworms (Oligochaeta, Annelida) in Field Studies. OECD Guidelines for the Testing of Chemicals, Section 2. OECD Publishing.

Oliveira, B.F., São-Pedro, V.A., Santos-Barrera, G., Penone, C. & Costa, G.C. (2017) AmphiBIO, a global database for amphibian ecological traits. Scientific Data 4, 170123.

Ortiz-Ceballos, C.E., AI Fragoso (2005) Influence of food quality, soil moisture and the earthworm *Pontoscolex corethrurus* on growth and reproduction of the tropical earthworm *Balanteodrilus pearseii*. Pedobiologia 49, 89–98.

Oskyrko, O., Mi, C., Meiri, S. & Du, W. (2024) ReptTraits: a comprehensive dataset of ecological traits in reptiles. Scientific Data 11, 243.

Palma, E., Vesk, P.A., White, M., Baumgartner, J.B. & Catford, J.A. (2021) Plant functional traits reflect different dimensions of species invasiveness. Ecology 102, e03317.

Pebesma, E. & Bivand, R. (2023) Spatial Data Science: With Applications in R. Chapman and Hall/CRC, New York.

Peigné, J., Cannavacuolo, M., Gautronneau, Y., Aveline, A., Giteau, J.L. & Cluzeau, D. (2009) Earthworm populations under different tillage systems in organic farming. Soil and Tillage Research 104, 207–214.

Pelosi, C., Barot, S., Capowiez, Y., Hedde, M. & Vandenbulcke, F. (2014) Pesticides and earthworms. A review. Agronomy for Sustainable Development 34, 199–228.

Pelosi, C., Bertrand, M., Makowski, D. & Roger-Estrade, J. (2008) WORMDYN: A model of *Lumbricus terrestris* population dynamics in agricultural fields. Ecological Modelling 218, 219–234.

Pelosi, C., Bertrand, M., Thenard, J. & Mougin, C. (2015) Earthworms in a 15 years agricultural trial. Applied Soil Ecology 88, 1–8.

Pelosi, C., Joimel, S. & Makowski, D. (2013) Searching for a more sensitive earthworm species to be used in pesticide homologation tests - A meta-analysis. Chemosphere 90, 895–900.

Pey, B., Nahmani, J., Auclerc, A., Capowiez, Y., Cluzeau, D., Cortet, J., Decaëns, T., Deharveng, L., Dubs, F., Joimel, S., Briard, C., Grumiaux, F., Laporte, M.-A., Pasquet, A., Pelosi, C., et al. (2014) Current use of and future needs for soil invertebrate functional traits in community ecology. Basic and Applied Ecology 15, 194–206.

Phillips, H.R.P. & Cameron, E.K. (2023) The GloWorm Project: Compiling regional level data to understand large-scale distributions of native and non-native species. Zenodo.

Phillips, H.R.P., Cameron, E.K., Eisenhauer, N., Burton, V.J., Ferlian, O., Jin, Y., Kanabar, S., Malladi, S., Murphy, R.E., Peter, A., Petrocelli, I., Ristok, C., Tyndall, K., Putten, W. van der & Beaumelle, L. (2024) Global changes and their environmental stressors have a significant impact on soil biodiversity—A meta-analysis. iScience 27.

Phillips, H.R.P., Guerra, C.A., Bartz, M.L.C., Briones, M.J.I., Brown, G., Crowther, T.W., Ferlian, O., Gongalsky, K.B., van den Hoogen, J., Krebs, J., Orgiazzi, A., Routh, D., Schwarz, B., Bach, E.M., Bennett, J.M., et al. (2019) Global distribution of earthworm diversity. Science 366, 480–485.

Phillipson, J. & Bolton, P. (1976) Respiratory metabolism of selected Lumbricidae. Oecologia 22, 135–152.

Plum, N. (2005) Terrestrial invertebrates in flooded grassland: A literature review. Wetlands 25, 721–737.

Posit team (2026) RStudio: Integrated development environment for R. manual, Posit Software, PBC, Boston, MA.

Potapov, A.M., Guerra, C.A., van den Hoogen, J., Babenko, A., Bellini, B.C., Berg, M.P., Chown, S.L., Deharveng, L., Kováč, Ľ., Kuznetsova, N.A., Ponge, J.-F., Potapov, M.B., Russell, D.J., Alexandre, D., Alatalo, J.M., et al. (2023) Globally invariant metabolism but density-diversity mismatch in springtails. Nature Communications 14, 674.

Prener, C., Grossenbacher, T., Zehr, A. & Stevens, J. (2025) biscale: Tools and Palettes for Bivariate Thematic Mapping. Https://cran.r-project.org/web/packages/biscale/index.html [accessed 18 May 2026].

Pyšek, P., Bacher, S., Kühn, I., Novoa, A., Catford, J.A., Hulme, P.E., Pergl, J., Richardson, D.M., Wilson, J.R.U. & Blackburn, T.M. (2020a) MAcroecological Framework for Invasive Aliens (MAFIA): disentangling large-scale context dependence in biological invasions. NeoBiota 62, 407–461.

Pyšek, P., Hulme, P.E., Simberloff, D., Bacher, S., Blackburn, T.M., Carlton, J.T., Dawson, W., Essl, F., Foxcroft, L.C., Genovesi, P., Jeschke, J.M., Kühn, I., Liebhold, A.M., Mandrak, N.E., Meyerson, L.A., et al. (2020b) Scientists’ warning on invasive alien species. Biological Reviews 95, 1511–1534.

R Core Team (2020) R: A language and environment for statistical computing. R Foundation for Statistical Computing, Vienna, Austria. Https://www.R-project.org/.

Rasmussen, L. & Holmstrup, M. (2002) Geographic variation of freeze-tolerance in the earthworm *Dendrobaena octaedra*. Journal of Comparative Physiology B-Biochemical Systemic and Environmental Physiology 172, 691–698.

Reinecke, A. & Kriel, J. (1981) Influence of temperature on the reproduction of the earthworm *Eisenia foetida* (Oligochaeta). South African Journal of Zoology 16, 96–100.

Reinecke, A., Viljoen, S. & Saayman, R. (1992) The suitability of *Eudrilus eugeniae*, *Perionyx excavatus* and *Eisenia foetida* (Oligochaeta) for vermicomposting in Southern Africa in terms of their temperature requirements. Soil Biology and Biochemistry 24, 1295–1307. Elsevier.

Richard, B., Decaëns, T., Rougerie, R., James, S.W., Porco, D. & Hebert, P.D.N. (2010) Re-integrating earthworm juveniles into soil biodiversity studies: species identification through DNA barcoding. Molecular Ecology Resources 10, 606–614.

Richardson, D.M. & Pyšek, P. (2006) Plant invasions: merging the concepts of species invasiveness and community invasibility. Progress in Physical Geography: Earth and Environment 30, 409–431.

Rillig, M.C., Ryo, M., Lehmann, A., Aguilar-Trigueros, C.A., Buchert, S., Wulf, A., Iwasaki, A., Roy, J. & Yang, G. (2019) The role of multiple global change factors in driving soil functions and microbial biodiversity. Science 366, 886–890.

Roani, R., Dudas, R.T., Demetrio, W.C., Lourenço, F.M.O., Ramos, G.A., Niemeyer, J.C., Tornisielo, V.L., Bonfleur, E.J., Ma, J.H.M., Bartz, M.L.C. & Brown, G.G. (2026) Pesticide residues and earthworm reproduction in 18 Brazilian soils. Environmental Toxicology and Chemistry 45, 613–631.

Rosner, A., Ballarin, L., Barnay-Verdier, S., Borisenko, I., Drago, L., Drobne, D., Concetta Eliso, M., Harbuzov, Z., Grimaldi, A., Guy-Haim, T., Karahan, A., Lynch, I., Giulia Lionetto, M., Martinez, P., Mehennaoui, K., et al. (2024) A broad-taxa approach as an important concept in ecotoxicological studies and pollution monitoring. Biological Reviews 99, 131–176.

Roy, H. E., Pauchard, A., Stoett, P., Renard Truong, T., Bacher, S., Galil, B. S., Hulme, P. E., Ikeda, T., Sankaran, K., McGeoch, M. A., Meyerson, L. A., Nuñez, M. A., Ordonez, A., Rahlao, S. J., Schwindt, E., Seebens, H., Sheppard, A. W., & Vandvik, V. (2023). IPBES Invasive Alien Species Assessment: Summary for Policymakers (Version 3). Zenodo. 10.5281/zenodo.10127924

Rundgren, S. (1975) Vertical distribution of lumbricids in Southern Sweden. Oikos 26, 299–306.

Sakai, A.K., Allendorf, F.W., Holt, J.S., Lodge, D.M., Molofsky, J., With, K.A., Baughman, S., Cabin, R.J., Cohen, J.E., Ellstrand, N.C., McCauley, D.E., O’Neil, P., Parker, I.M., Thompson, J.N. & Weller, S.G. (2001) The population biology of invasive species. Annual Review of Ecology, Evolution, and Systematics 32, 305–332.

Salyer, A., Bennett, G.W. & Buczkowski, G.A. (2014) Odorous house ants (*Tapinoma sessile*) as back-seat drivers of localized ant decline in urban habitats. PLoS ONE 9, e113878.

Sandmann, D., Scheu, S. & Potapov, A. (2019) Ecotaxonomy: Linking taxa with traits and integrating taxonomical and ecological research. Biodiversity Information Science and Standards 3, e37146.

Scott-Fordsmand, J., Weeks, J. & Hopkin, S. (1998) Toxicity of nickel to the earthworm and the applicability of the neutral red retention assay. Ecotoxicology 7, 291–295.

Seebens, H., Blackburn, T.M., Dyer, E.E., Genovesi, P., Hulme, P.E., Jeschke, J.M., Pagad, S., Pyšek, P., Winter, M., Arianoutsou, M., Bacher, S., Blasius, B., Brundu, G., Capinha, C., Celesti-Grapow, L., et al. (2017) No saturation in the accumulation of alien species worldwide. Nature Communications 8, 14435.

Simberloff, D. (2009) The Role of Propagule Pressure in Biological Invasions. Annual Review of Ecology, Evolution, and Systematics 40, 81–102.

Sinclair, J.S., Lockwood, J.L., Hasnain, S., Cassey, P. & Arnott, S.E. (2020) A framework for predicting which non-native individuals and species will enter, survive, and exit human-mediated transport. Biological Invasions 22, 217–231.

Sinha, R.K., Herat, S., Agarwal, S., Asadi, R. & Carretero, E. (2002) Vermiculture and waste management: Study of action of earthworms *Eisinia foetida*, *Eudrilus euginae* and *Perionyx excavatus* on biodegradation of some community wastes in India and Australia. The Environmentalist 22, 261–268.

Sjoberg, D. (2026) davidsjoberg/ggsankey. R,. Https://github.com/davidsjoberg/ggsankey [accessed 18 May 2026].

Smith, C.C. & Fretwell, S.D. (1974) The optimal balance between size and number of offspring. The American Naturalist 108, 499–506.

Smith, R.G., McSwiney, C.P., Grandy, A.S., Suwanwaree, P., Snider, R.M. & Robertson, G.P. (2008) Diversity and abundance of earthworms across an agricultural land-use intensity gradient. Soil and Tillage Research 100, 83–88.

Sol, D., Maspons, J., Vall-llosera, M., Bartomeus, I., García-Peña, G.E., Piñol, J. & Freckleton, R.P. (2012) Unraveling the life history of successful invaders. Science 337, 580–583.

Stevenson, R.D. (1985) Body size and limits to the daily range of body temperature in terrestrial ectotherms. The American Naturalist 125, 102–117.

Svendsen, C., Hankard, P.K., Lister, L.J., Fishwick, S.K., Jonker, M.J. & Spurgeon, D.J. (2007) Effect of temperature and season on reproduction, neutral red retention and metallothionein responses of earthworms exposed to metals in field soils. Environmental Pollution 147, 83–93.

Terhivuo, J. & Saura, A. (2006) Dispersal and clonal diversity of North-European parthenogenetic earthworms. Biological Invasions 8, 1205–1218.

Tobias, J.A., Sheard, C., Pigot, A.L., Devenish, A.J.M., Yang, J., Sayol, F., Neate-Clegg, M.H.C., Alioravainen, N., Weeks, T.L., Barber, R.A., Walkden, P.A., MacGregor, H.E.A., Jones, S.E.I., Vincent, C., Phillips, A.G., et al. (2022) AVONET: morphological, ecological and geographical data for all birds. Ecology Letters 25, 581–597.

Ulrich, S., Tischer, S., Hofmann, B. & Christen, O. (2010) Biological soil properties in a long-term tillage trial in Germany. Journal of Plant Nutrition and Soil Science 173, 483–489.

Uvarov, A. & Scheu, S. (2004) Effects of temperature regime on the respiratory activity of developmental stages of *Lumbricus rubellus* (Lumbricidae). Pedobiologia 48, 365–371.

Van Gestel, C.A.M. & Weeks, J.M. (2004) Recommendations of the 3rd International Workshop on Earthworm Ecotoxicology, Aarhus, Denmark, August 2001. Ecotoxicology and Environmental Safety 57, 100–105.

Vanadzina, K., Nagahawatte, K., Gérard, S., Cameron, E., Brown, G., Reynolds, J., Rice, A. & Phillips, H. (2025) Systematic evidence map of research on life-history traits in native and introduced earthworms. OSF.

Vasseur, P. & Bonnard, M. (2014) Ecogenotoxicology in earthworms: A review. Current Zoology 60, 255–272.

Walsh, B.S., Parratt, S.R., Hoffmann, A.A., Atkinson, D., Snook, R.R., Bretman, A. & Price, T.A.R. (2019) The impact of climate change on fertility. Trends in Ecology & Evolution 34, 249–259.

Walther, G.-R., Roques, A., Hulme, P.E., Sykes, M.T., Pyšek, P., Kühn, I., Zobel, M., Bacher, S., Botta-Dukát, Z., Bugmann, H., Czúcz, B., Dauber, J., Hickler, T., Jarošík, V., Kenis, M., et al. (2009) Alien species in a warmer world: risks and opportunities. Trends in Ecology & Evolution 24, 686–693.

Wickham, H. (2023) _stringr: Simple, Consistent Wrappers for Common String Operations. Https://CRAN.R-project.org/package=stringr [accessed 13 May 2026].

Wickham, H., Averick, M., Bryan, J., Chang, W., McGowan, L.D., François, R., Grolemund, G., Hayes, A., Henry, L., Hester, J., Kuhn, M., Pedersen, T.L., Miller, E., Bache, S.M., Müller, K., et al. (2019) Welcome to the Tidyverse. Journal of Open Source Software 4, 1686.

Winsome, T., Epstein, L., Hendrix, P. & Horwath, W. (2006) Competitive interactions between native and exotic earthworm species as influenced by habitat quality in a California grassland. Applied Soil Ecology 32, 38–53.

Yasmin, S. & D’Souza, D. (2007) Effect of pesticides on the reproductive output of *Eisenia fetida*. Bulletin of Environmental Contamination and Toxicology 79, 529–532.

Zhang, C., Wright, I.J., Nielsen, U.N., Geisen, S. & Liu, M. (2024) Linking nematodes and ecosystem function: a trait-based framework. Trends in Ecology & Evolution 39, 644–653.

Zhang, L., Shen, W., Fang, Z., Liu, L., Jia, R., Liang, J. & Liu, B. (2025) Multigenerational effects of cultivating transgenic maize straw on earthworms: A combined laboratory and field experiment. Ecotoxicology and Environmental Safety 291.

Zhang, M., Jouquet, P., Dai, J., Xiao, L., Du, Y., Liu, K., Motelica-Heino, M., Lavelle, P., Zhong, H. & Zhang, C. (2022) Assessment of bioremediation potential of metal contaminated soils (Cu, Cd, Pb and Zn) by earthworms from their tolerance, accumulation and impact on metal activation and soil quality: A case study in South China. Science of The Total Environment 820, 152834.

